# Breastmilk IgG engages the neonatal immune system to instruct host-microbiota mutualism

**DOI:** 10.1101/2024.08.23.609293

**Authors:** Meera K. Shenoy, Diane Rico, Shannon Gordon, Luke Milburn, Jeanette Schwensen, Madelyn Cabàn, Meghan A. Koch

## Abstract

Maternal antibodies fundamentally regulate infant immunity to the developing gut microbiota, yet the mechanisms underlying this process remain elusive. Here, we show that maternal IgG, ingested in the first week of life, functions to restrain microbiota-dependent adaptive immune responses and reduce offspring susceptibility to intestinal inflammation weeks later, following weaning. To exert these functions, efficient binding of IgG to gut bacterial antigens and engagement of Fc and complement dependent effector functions in offspring was required. These discoveries reveal a novel mechanism wherein maternal IgG engages the offspring immune system to calibrate responses to gut microbes. This mode of maternal immune instruction may provide adaptability to developmental shifts in microbiota necessary for establishing host-microbiota mutualism and limiting susceptibility to inflammatory disease.

**One sentence abstract:** Ingestion of maternal IgG during a discrete postnatal window calibrates neonatal immunity to the gut microbiota.

## Introduction

A fundamental function of the immune system is to promote mutualism between the host and resident microbiota. In the gut, this process enables the host to harness numerous benefits from microbes, including expanded nutrient assimilation, immune education, and colonization resistance to pathogens, while mitigating risks such as toxin exposure, chronic inflammation, and pathobiont invasion (1–3). Establishing this balance is complicated by ongoing development of neonatal immunity, coupled with developmental- and diet-directed dynamic shifts in microbial composition. Nonetheless, studies linking perturbations in infant microbiota exposure or composition with increased risk of developing metabolic and/or inflammatory diseases underscore the importance of this process (4–9). Ingested immediately post-birth, breastmilk is poised to regulate nascent host-microbiota interactions. Along with nutrients, breastmilk provides offspring with live bacteria, maternal immune cells, and immunomodulatory molecules including cytokines and immunoglobulins (10, 11).

Breastmilk contains several antibody isotypes including IgA and IgG, which readily bind microbes in the infant gut and help shape assembly of the microbiome, promote barrier function, restrain inflammatory responses to commensal microbes, and guide the abundance of mucosal T cells (1–7). The prevailing model of maternal antibody-mediated immune instruction is largely based on studies of IgA regulation of enteric pathogens or certain commensal strains in the intestines of adult mice (8–14). However, this model does not account for substantial differences in barrier physiology, microbiota dynamics, and luminal antibody composition (e.g., IgG) in the infant gut (7, 15–18). As such, the mechanisms by underlying antibody function at the maternal-offspring interface remain unknown. Here, we developed a murine model enabling precise control over the timing and composition of breastmilk antibodies, the microbiota, and the infant immune system to explore the possibility that breastmilk antibodies operate via unique mechanisms to tune offspring immunity to the emerging microbiota.

## Results

### Early life acquisition of breastmilk antibodies restrains mucosal immune dysregulation during the weaning transition

We previously reported that breastmilk antibodies regulate mucosal adaptive immunity in offspring, wherein wild-type C57Bl/6 (B6) offspring reared by μMT^-/-^ dams (that lack mature B cells and thus do not produce antibodies) exhibit heightened Germinal Center (GC) T follicular helper (Tfh) and B cells in gut-draining mesenteric lymph nodes (mLN) and Peyer’s patches (PP) (7). Accumulating evidence indicates that establishing beneficial immunity to microbiota relies on a series of coordinated events occurring during distinct developmental phases (19). To determine whether breastmilk antibody transfer during a specific postnatal window was required to regulate mucosal immunity, we bred B cell-sufficient μMT^+/-^ and -deficient μMT^-/-^ dams to B6 sires and cross-fostered their offspring at different ages (**Fig 1A**). To ensure maximum homogenization of the microbiota between birth mothers, offspring, and foster mothers, littermate, cohoused μMT^+/-^ and μMT^-/-^ dams were briefly separated for breeding with sires, and again prior to parturition and through offspring weaning.

**Figure 1.**
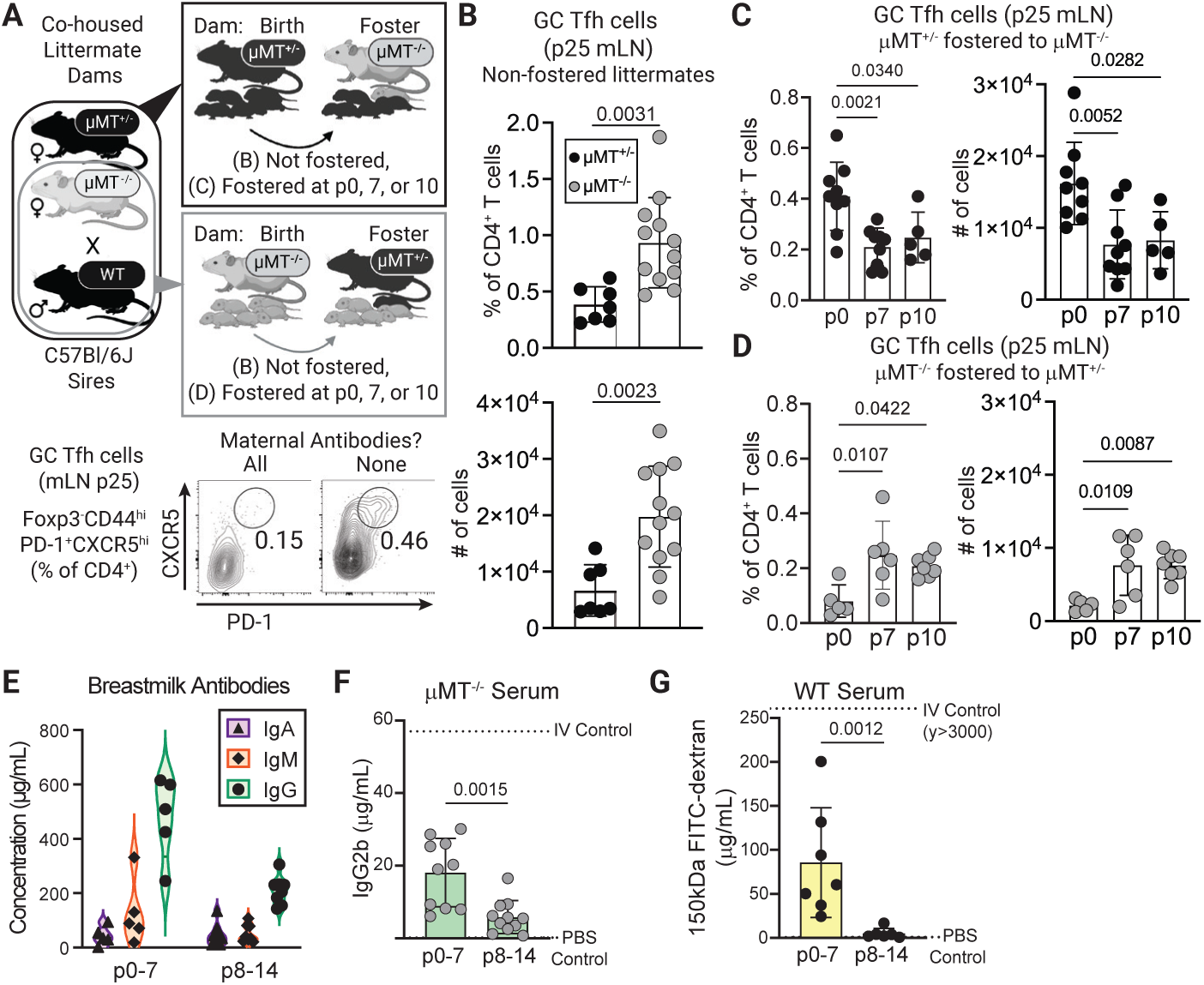
Acquisition of breastmilk antibodies in early life prevents immune dysregulation following weaning. (**A**) (Top Schema) Timed breedings were performed using co-housed littermate μMT^+/-^ and μMT^-/-^ dams and B6 sires. Offspring were maintained with their birth mother or cross-fostered at postnatal day 0, 7, or 10, and analyzed at p25. (Bottom) Representative flow cytometric analysis of GC Tfh cells (CD4^+^Foxp3^-^CD44^hi^PD-1^+^CXCR5^+^) isolated from the mLN of p25 offspring born to and reared by the indicated dams. (**B**) Proportions (top) and numbers (bottom) of mLN GC Tfh cells in p25 offspring born to and reared by the indicated dams. (**C**) Proportions (left) and numbers (right) of mLN GC Tfh cells in p25 offspring born to μMT^+/-^ dams and fostered to μMT^-/-^ dams at the indicated postnatal timepoints. (**D**) Similar to (C) comparing μMT^-/-^ born offspring fostered to μMT^+/-^ dams. (**E**) Breastmilk IgA, IgM, and IgG titers of B6 dams sampled at the indicated time periods post-parturition. All dams were 12-16 weeks of age. (**F**) Serum IgG2b titers in μMT^-/-^ pups of the indicated age 2 hours post gavage with 10mg/kg monoclonal IgG2b antibodies. Lower and upper dotted lines indicate IgG2b concentrations in mice gavaged with PBS or injected i.v. with IgG2b, respectively. (**G**) Serum FITC-dextran (FITC-dex) titers in wild-type pups of the indicated age 2 hours post gavage with 10mg/kg FITC-dex (150kDa). Lower and upper dotted lines indicate FITC-dex concentrations in mice gavaged with PBS or injected i.v. with FITC-dex, respectively. Error bars indicate the mean ±SD; symbols represent individual mice. Data are representative of two to four independent experiments with ≥ 5 mice per group. Statistical significance was determined using one-way ANOVA and Tukey post-hoc tests, except (B), (F), and (G), wherein an unpaired two-tailed Student’s *t* test was used.

Mirroring our prior findings (7), acquisition of breastmilk antibodies during the entire post-weaning period prevented mucosal immune dysregulation. Compared with control pups reared by μMT^+/-^ dams, those nursed by μMT^-/-^ dams mounted elevated Tfh (CD4^+^Foxp3^-^CD44^+^PD-1^+^CXCR5^hi^, **Fig S1A**) and GC B (CD19^+^IgD^lo^B220^+^PNA^+^GL7^+^, **Fig S1B**) cell responses in the mLN and PP at postnatal day 25 (p25; (**Fig 1B-D, Fig S1C-H**)). Strikingly, progeny of μMT^+/-^ dams fostered onto antibody-deficient μMT^-/-^ dams after the first week of life (p7 or later) harbored reduced frequencies and numbers of Tfh and GC B cells compared to pups fostered at birth (p0), indicating that breastmilk antibody ingestion during the first week of life is sufficient to prevent adaptive immune dysregulation during the weaning transition (**Fig 1C, Fig S1G**). Corroborating these findings, μMT^-/-^ dam-born pups fostered onto antibody-sufficient μMT^+/-^ dams at p0 did not exhibit immune dysregulation, whereas elevated mucosal Tfh and GC B cell responses were apparent in littermate counterparts fostered after p7 (**Fig 1D, Fig S1H**).

### Kinetics of neonatal antibody consumption and gut permeability

To start to understand how breastmilk antibodies regulate intestinal homeostasis, we assessed breastmilk consumption and composition over time (**Fig S2A**). Expectedly, breastmilk intake increased as a function of age (**Fig S2B**), and the total immunoglobulin concentration in milk (the sum of IgG, IgA, and IgM; IgE not detected) was largely stable during the first two weeks of lactation. Interestingly, milk from dams 0-7 days postpartum contained significantly more IgG than those sampled later (**Fig 1E**). Averaging milk intake and antibody concentrations, we estimate pups in our mouse colony consume 244 μg/day of IgG and 17 μg/day IgA in the first week of life (**Fig S2C**).

The developing gastrointestinal tract exhibits physiological changes characterized by alterations in antimicrobial peptides, mucus, nutrient transporters, and barrier permeability (20, 21). In healthy human infants, gut permeability to small sugars is increased during the first week of life (22), and one study reported translocation of IgA dimers (∼300kDa) in term infants orally administered IgA within 24 hours post-birth (23). Thus, we assessed whether the window of breastmilk antibody acquisition required to regulate offspring immunity correlated with gut barrier permeability to ingested IgG. To directly evaluate IgG translocation in the absence of cognate antigens or endogenous maternal- or host-derived IgG, we orally administered anti-ovalbumin IgG2b to μMT^-/-^ offspring of μMT^-/-^ dams. Two hours after treatment, we detected higher levels of serum IgG2b in p0-7 offspring compared to older pups (**Fig 1F**). We observed similar results in B6 offspring fed 150kDa Fluorescein-conjugated dextran (similar in size to IgG) (**Fig 1G**), ruling out the possibility that the observed increase in barrier permeability resulted from developmental defects in μMT^-/-^ mice. Thus, the mouse gut is more permeable to luminal substrates during the first week of life: the same window during which maternal antibodies must be ingested to establish intestinal homeostasis.

### Oral IgG is sufficient to prevent mucosal immune dysregulation in offspring

To define the maternal antibody isotypes required to prevent immune dysregulation, we fractionated mouse milk IgG from IgA and other proteins (**Fig 2A**, **Fig S3A**) and fed littermate, maternal-antibody deficient offspring 0.5 μg of milk IgG (mIgG), 1μg milk IgA (which contained other proteins; mIgA+prot), or 1μg BSA daily from p0 to p7. At p25, neonates fed mIgG exhibited reduced mucosal Tfh and GC B cell responses compared to pups administered BSA or IgA plus other milk proteins (**Fig 2B**, **Fig S3B-C**). We obtained similar results using purified serum IgG (sIgG), wherein administration of sIgG, but not serum depleted of IgG (sΔIgG) resulted in significantly fewer Tfh and GC B cells in p25 progeny of μMT^-/-^ dams x B6 sires (**Fig 2C, Fig S3D-E**). Importantly, this ‘add-back’ approach allows a direct comparison of co-housed littermates born to and reared by the same mother, thus avoiding potential confounding variables introduced by maternal microbiota, antibody concentrations, or housing conditions (24). Indeed, though the magnitude of the immune dysregulation varied between litters, mLN Tfh cell responses were consistently lower in pups fed sIgG compared to littermate controls given sΔIgG (**Fig 2D, Fig S3F-G**). The ability of IgG to prevent mucosal immune dysregulation indicates that the absence of antibodies, rather than an unrelated perturbation in the milk or behavior of μMT^-/-^ mothers, triggers aberrant intestinal immunity. Dose-response experiments revealed that as little as 1 μg/day sIgG during p0-7 was sufficient to fully restrain mesenteric Tfh and GC B cell responses in p25 offspring of μMT^-/-^ dams and B6 sires (**Fig 2E, Fig S3H**). As mouse serum is more plentiful than milk, and milk lipoproteins impede purification of IgG, we used purified sIgG for the remaining studies.

**Figure 2.**
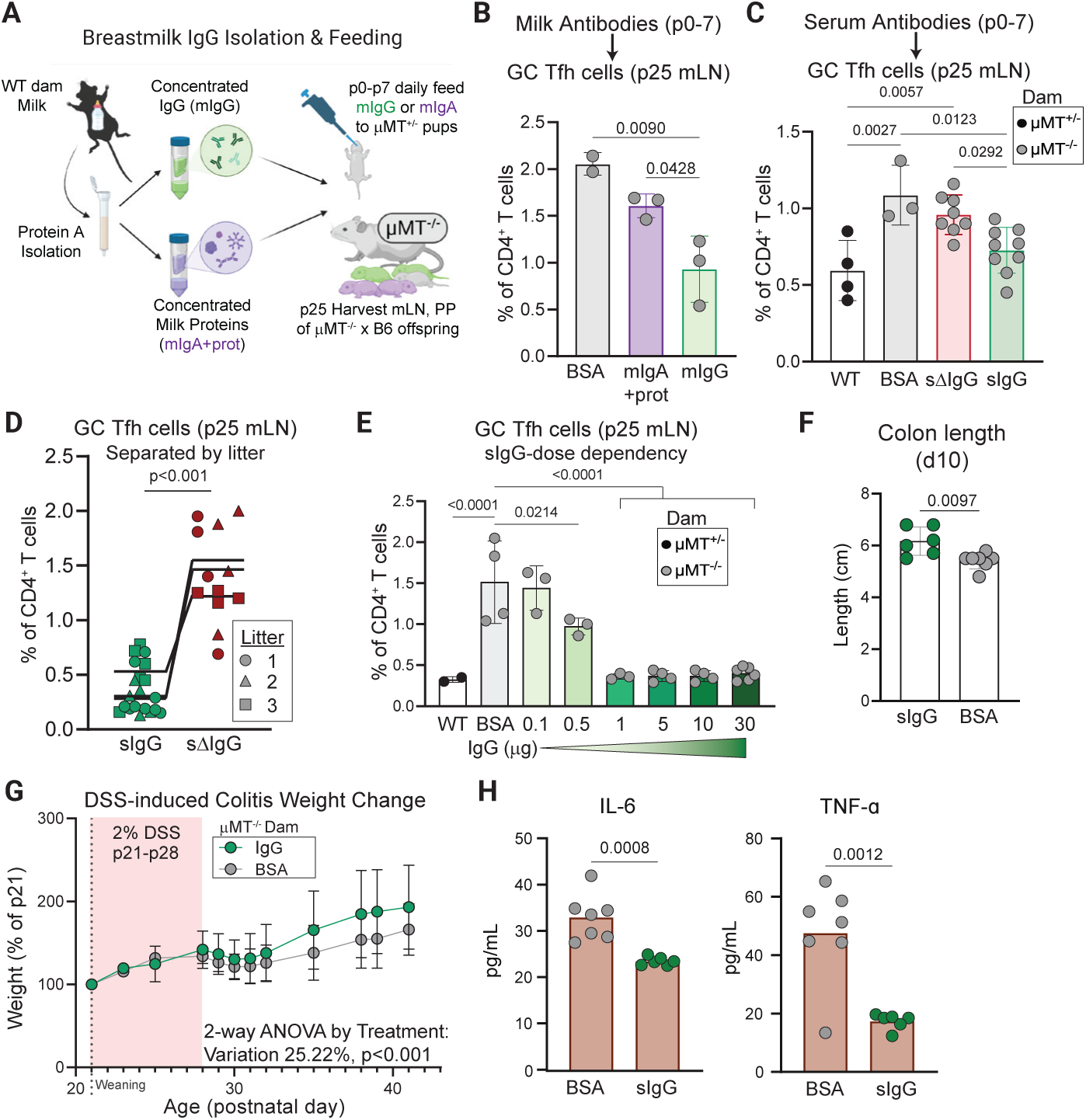
Oral IgG is sufficient to regulate immunity to the microbiota in offspring lacking maternal antibodies. (**A**) Milk from B6 dams was enriched into IgG+ and IgG-fractions, concentrated, and fed to offspring of μMT^-/-^ dams and B6 sires during the first week of life. Control pups were fed BSA. (**B**) Proportions of mLN GC Tfh cells in p25 offspring fed 0.5μg milk (m)IgG, 1μg mIgA (+ other proteins), or 1μg BSA daily from p0-p7. (**C**) Similar to (B) comparing proportions of mLN GC Tfh cells in p25 offspring fed 10μg serum purified IgG (sIgG), IgG-depleted serum (sΔIgG) or BSA daily during p0-p7. (**D**) Similar to (B), comparing proportions of mLN GC Tfh cells in littermate, co-housed pups given 10μg sIgG or sΔIgG between p0-p7. Littermate animals indicated by symbols. (**E**) Similar to (B), comparing p25 mLN GC Tfh responses in offspring fed the indicated concentrations of sIgG or BSA between p0-p7. (**F**) Offspring of μMT^-/-^ dams and B6 sires were fed purified 10μg sIgG or BSA during the first week of life, weaned and cohoused with offspring of μMT^+/-^ dams and B6 sires at p20 and given 2% DSS in their drinking water for 1 week. Graph depicts colon lengths of the indicated offspring 10 days post DSS administration. (**G**) Median percentage of original weight (±SD) of the indicated offspring following DSS administration as described in (F). Variation across mice was statistically significant by treatment group, *p<0.0001,* by 2-way ANOVA. (**H**) Serum titers of the indicated offspring 10 days following DSS administration as described in (F). For (**B** to **F,** and **H**) error bars indicate the mean ±SD; symbols represent individual mice. Data are representative of two - four independent experiments with ≥ 3 mice per group. Statistical significance was determined using one-way ANOVA and Tukey post-hoc tests, except for (**D, F** and **H**), wherein an unpaired two-tailed Student’s *t* test was used. For (**G**) symbols indicate the mean and error bars indicate ±SD. Data in are representative of three independent experiments with ≥ 8 mice per group.

In addition to regulating homeostatic immunity, maternal antibodies can promote health during inflammatory contexts (7, 18). We found that provision of oral IgG in early life protected mice from experimental colitis induced during the weaning transition. Compared with BSA-treated littermates, sIgG supplementation of progeny of μMT^-/-^ dams and B6 sires resulted in greater weight gain, longer colon lengths, and reduced inflammatory cytokine levels induced by administration of DSS (**Fig 2F-H)**.

### An intact gut microbiota is required to trigger mucosal immune dysregulation in offspring

The selective elevation of Tfh and GC B cell responses at mucosal (e.g., mLN and PP) but not systemic sites (e.g., spleen) (7), suggests that gut microbes trigger immune dysregulation in offspring lacking maternal antibodies. To test this hypothesis, we analyzed maternal antibody-sufficient and -deficient offspring housed under germ-free (GF) conditions. At p25, neonates harbored equivalent, low frequencies and numbers of mucosal Tfh and GC B cells, regardless of maternal antibody acquisition (**Fig 3A, Fig S4A**). Next, we depleted the microbiota in specific pathogen free (SPF) mice by administering broad spectrum antibiotics (ampicillin, vancomycin, neomycin, and metronidazole; AVNM (25)) during the weaning transition (**Fig 3B**). At p25, AVNM-treated pups lacking maternal antibodies harbored significantly reduced frequencies and numbers of mucosal Tfh and GC B cell responses compared to their non-AVNM-treated littermates (**Fig 3C, Fig S4B, C**). Thus, the gut microbiota triggers immune dysregulation in offspring lacking maternal antibodies. Interestingly, weaning pups earlier (p18) or later (p25) than the standard of p21 resulted in similar immune dysregulation, indicating that this microbiota-elicited response is not controlled by the cessation of breastfeeding (**Fig 3D, Fig S4D, E**).

**Figure 3.**
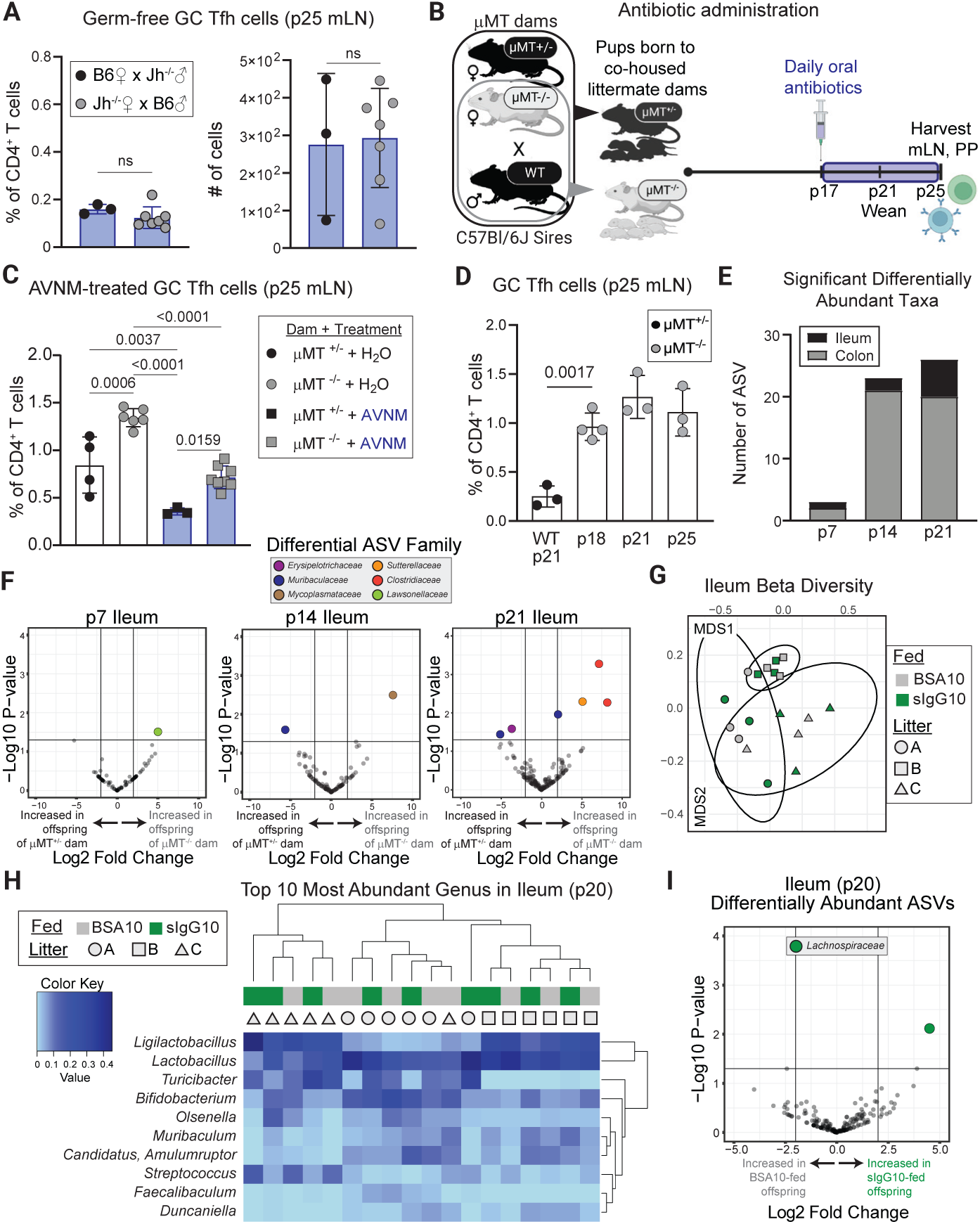
Impact and assembly of the microbiota in maternal antibody deficient offspring. (**A**). Proportions (left) and numbers (right) of mLN GC Tfh cells in germ-free p25 offspring born to and reared by the indicated dams bred with the indicated sires. (**B**) Timed breedings were performed using co-housed littermate μMT^+/-^ and μMT^-/-^ dams and B6 sires. Offspring were orally administered antibiotics (AVNM) or water daily between p17-p24. (**C**) Proportions of mLN GC Tfh cells in p25 offspring born to the indicated dams and given the indicated treatments. (**D**) Proportions of mLN GC Tfh cells in p25 offspring born to the indicated dams and weaned at the indicated postnatal timepoints. (**E**) Number of significant differentially abundant taxa (sequence variants) between maternal antibody-sufficient and -deficient offspring in the indicated intestinal sites at the indicated postnatal timepoints. (**F**) Volcano plot of ileal microbiota taxa (sequence variants) at the indicated postnatal timepoints of mice described in (E). (**G**) Non-metric multidimensional scaling plot displaying the Bray-Curtis distances between ileal bacteria from p20 littermate offspring of μMT^-/-^ dams and B6 sires fed 10 μg sIgG or BSA in the first week of life. PERMANOVA analysis: Litter p=0.001, R^2^=0.633 and IgG feeding group p=0.746, R^2^=0.01. (**H**) Heatmap of genus-level ileal taxa (rows) isolated from individual p20 offspring (columns) described in (G) and ordered by complete linkage clustering. Feed and litter status of each mouse is indicated on the top of each heatmap by colors and symbols, respectively. (**I**) Volcano plot of p20 ileal microbiota species (sequence variants) in mice described in (G). The taxonomic family of the single significantly altered taxa is indicated. For (**A** to **D**) Error bars indicate the mean ±SD; symbols represent individual mice. Data are representative of three independent experiments with ≥ 3 mice per group. Statistical significance was determined using one-way ANOVA and Tukey post-hoc tests. For (**E** and **F**) Data are combined from 2 independent experimental cohorts with samples harvested greater than one year apart totaling n≥10/group. For (**G** to **I**) data are generated from 3 separate litters with n=9 per group. Symbols in (**G** and **H**) and columns in (**I**) represent individual mice.

### The impact of maternal antibodies on the assembly of the postnatal microbiome

The requirement of the microbiota to elicit immune dysregulation coupled with the established role of antibodies in regulating gut microbiota diversity and composition (5, 26–28) prompted us to explore whether enhanced Tfh responses correlated with alterations in microbiome assembly in maternal antibody-deficient offspring. 16S rRNA sequencing of ileal and colonic contents (**Fig S5A**) did not reveal substantial differences in the overall diversity of microbes (alpha diversity) nor community composition (beta diversity) between age-matched offspring of μMT^+/-^ or μMT^-/-^ dams and B6 sires (**Fig S5B-E**). However, the abundance of a small subset of taxa differed significantly between p7 offspring of μMT^+/-^ and μMT^-/-^ dams, and this variation increased with age (**Fig 3E, F, Fig S5F-H)**. Thus, the absence of all breastmilk antibodies correlates with minor alterations in the assembly of the postnatal microbiome. Analysis of p20 progeny of μMT^-/-^ dams and B6 sires that received sIgG or BSA during the first week of life similarly failed to reveal significant differences in alpha or beta diversity as a function of feeding (**Fig 3G, Fig S6A-C**). Instead, litter was a driving factor (**Fig 3H, Fig S6D**). Finally, the provision of IgG resulted in the differential abundance of only a single ileal taxon and two colonic taxa (**Fig 3I, S6E**), which were distinct from those identified in p21 offspring of μMT^+/-^ and μMT^-/-^ dams. Taken together with results in **Fig 2D** and **Fig S3F-G**, these data demonstrate that the ability of early life IgG to regulate neonatal immunity is robust to natural microbiome variation between litters. Additionally, the similarity in microbiota community structure between IgG and BSA fed littermates indicates that early life IgG does not regulate neonatal immunity by altering the composition of the intestinal microbiome.

### Early life IgG-microbe immune complexes prevent intestinal immune dysregulation

We previously showed that acquisition of maternal antibodies via breastfeeding, rather than placental transfer, was required to prevent Tfh-driven immune dysregulation (**Fig 1B**-**D** and (7)). Serum analysis of p7 offspring of μMT^+/-^ or μMT^-/-^ dams and B6 sires cross-fostered at birth revealed that a substantial amount of maternal IgG is obtained via breastfeeding as opposed to *in utero* transmission (**Fig 4A, S7A**). Strikingly, maternal antibody-deficient pups fed sIgG harbored even lower serum IgG titers compared to offspring that only received maternal antibodies *in utero*, ruling out the possibility that the immune dysregulation observed in pups nursed by antibody deficient dams resulted from insufficient titers of circulating IgG in early life (**Fig 4A, S7A**). As administration of sIgG is sufficient to reduce mucosal Tfh responses at p25, these findings indicate that oral acquisition, rather than circulating postnatal levels, underlies the ability of maternal IgG to calibrate offspring intestinal immunity.

**Figure 4.**
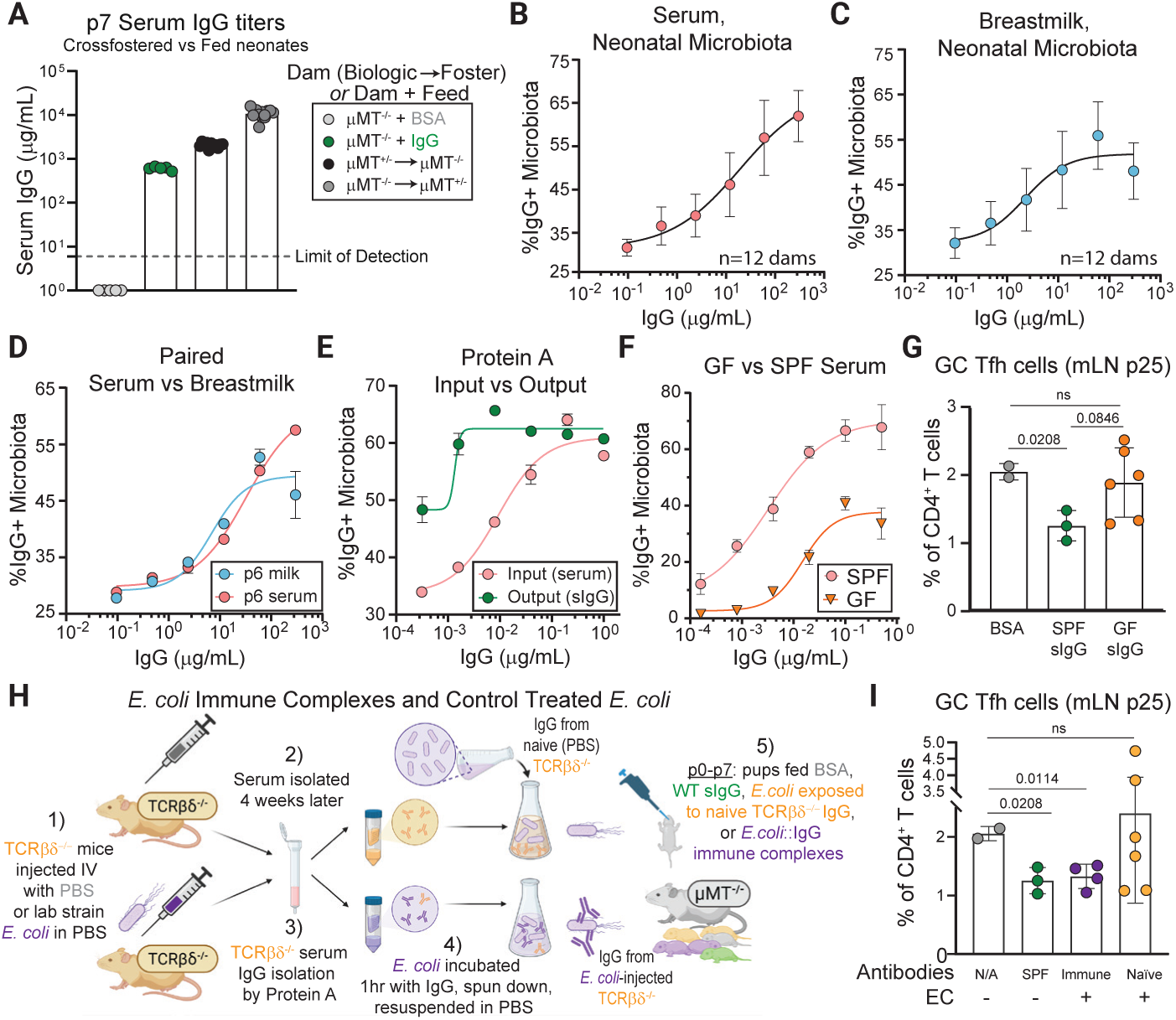
Efficient binding to mucosal antigens is required for early life IgG to restrain mucosal Tfh responses. (**A**) Serum IgG titers of p7 offspring of the indicated dams that were cross-fostered or fed proteins as indicated. (**B** to **D**) mFLOW analysis of the frequency of SYBR+ bacteria bound by IgG following incubation with sera (B), or milk (C), of multiple B6 donors (n>12), and matched sera and milk of a single donor (D). Microbes were isolated from the small intestine and colon of p6 μMT^-/-^ neonates. (**E**) Similar to (B) but comparing IgG-bound bacteria following incubation with serum IgG (input) or Protein A purified IgG as indicated. (**F**) Similar to (B) but comparing IgG-bound bacteria following incubation with sera of SPF or germ-free (GF) animals. (**G**) Proportions of mLN GC Tfh cells in p25 offspring of μMT^-/-^ dams and B6 sires fed 0.5μg sIgG purified from SPF or GF mice, or BSA, as indicated. (**H**) Serum was collected from TCRβδ^-/-^ mice 4 weeks after i.v. injection of *E. coli* or PBS and IgG was purified via Protein A fractionation. Timed breedings were performed using co-housed littermate μMT^+/-^ and μMT^-/-^ dams and B6 sires and resulting offspring were fed sIgG, BSA, or *E. coli* pre-incubated with IgG from either *E.coli* or PBS immunized mice daily in the first week of life. (**I**) Proportions of mLN GC Tfh cells in p25 offspring of μMT^-/-^ dams and B6 sires fed the indicated proteins +/- bacteria. For (**B** to **F**) Graphs depict the relative median ±SD. For (**A, G**, and **I**) error bars indicate the mean±SD; symbols represent individual mice. Data are representative of two to four independent experiments with ≥2 mice per group. Statistical significance was determined using one-way ANOVA and Tukey post-hoc tests.

These observations prompted us to hypothesize that maternal IgG must bind to neonatal gut microbes to establish host-microbiota mutualism. We first assessed the antibody binding profiles of milk and serum using microbiota-flow cytometry (mFLOW) (7). To avoid pre-coating of infant bacteria by maternal antibodies, we used gut contents from μMT^-/-^ pups (p6). IgG antibodies from either milk or serum bound a substantial fraction of the neonatal microbiota (**Fig 4B-D**). Interestingly, when matched for concentration, milk IgG tended to bind a larger fraction of neonatal gut bacteria, while paired serum IgG bound a larger proportion of adult gut microbes (derived from 3-month-old μMT^-/-^ mice) (**Fig 4D, Fig S7B**). Additionally, we confirmed that fractionated IgG used for feeding studies retained the ability to bind to neonatal gut microbes (**Fig 4E**).

Next, we tested whether efficient binding to intestinal microbes was required for oral IgG to prevent neonatal mucosal immune dysregulation. Using serum IgG from germ-free mice (GF-sIgG), which harbor reduced titers of microbiota-reactive antibodies (**Fig 4F, Fig S7C** (7, 29)), we fed progeny of μMT^-/-^ dams and B6 sires 1 μg/day GF-sIgG, SPF-sIgG, or BSA between p0-7. Though normalized by concentration, GF-sIgG was unable to restrain mucosal immune dysregulation as evidenced by heightened mucosal Tfh responses these animals compared to pups fed SPF-sIgG (**Fig 4G, Fig S7D, E**).

To complement these findings, we asked whether antibody-bacteria immune complexes were sufficient to prevent immune dysregulation in pups that did not receive breastmilk antibodies. To recapitulate the homeostatic T-independent, anti-commensal IgG responses observed in mice (7), we immunized TCRβẟ^-/-^ mice with *E. coli* strain BL21 (EC). Importantly, while *E. coli* is a normal constituent of the early life microbiota, naïve mice in our colony do not harbor antibodies that bind to this strain. Immunization elicited robust EC-specific IgG responses, which we used to generate immune complexes that were stable for up to one week (**Fig S7F**).

Strikingly, administration of EC immune complexes (Immune-EC) during the first week of life was sufficient to limit p25 mucosal Tfh responses to the level observed in offspring fed sIgG (**Fig 4H, I** and **Fig S7G, H**). In contrast, Tfh responses were equivalently elevated in mice given EC incubated with sera from naïve mice (naïve-EC) or BSA, indicating that early life encounter of EC alone did not prevent immune dysregulation. Taken together with our findings using GF-sIgG, these data indicate that maternal IgG must bind microbes in the infant gut and form immune complexes to tune neonatal immunity to the developing microbiota.

### Maternal IgG engages neonatal IgG sensors to restrain microbiota-dependent adaptive immunity

The Fc domain of mouse IgG, but not IgA, immune complexes allows them to shape immune responses by engaging IgG-sensing systems, such as Fc receptors (FcRs) and complement activation (30). To define the molecular basis by which maternal IgG restrains neonatal immune dysregulation, we fostered FcRγ^-/-^ offspring, which lack the common FcRγ chain required for the expression and signaling of activating FcRs (FcγRI, FcγRIII, and FcγRIV (31)) to antibody-deficient μMT^-/-^ dams at p0 and fed them sIgG or BSA daily for the first week of life (**Fig 5A**). Administration of sIgG resulted in a significant decrease in the frequency, but not number, of mLN Tfh and GC B cells in p25 FcRγ^-/-^ offspring (**Fig 5B, Fig S8A-B**). We observed similar, partial results using C1qa^-/-^ offspring, which are deficient in antibody-dependent complement activation (32). Specifically, IgG ingestion resulted in a slight reduction in mucosal Tfh and GC B cell responses in C1qa^-/-^ mice at p25 compared with BSA-fed progeny; however, this effect was less pronounced than that observed in C1qa-sufficient (control) progeny (**Fig 5C**, **Fig S8C-D**). Antibody dependent complement activation was confirmed via elevated intestinal C5a levels (a product of complement activation) in p7 C1qa-sufficient progeny given sIgG, but not BSA **(Fig S8E).**

**Figure 5.**
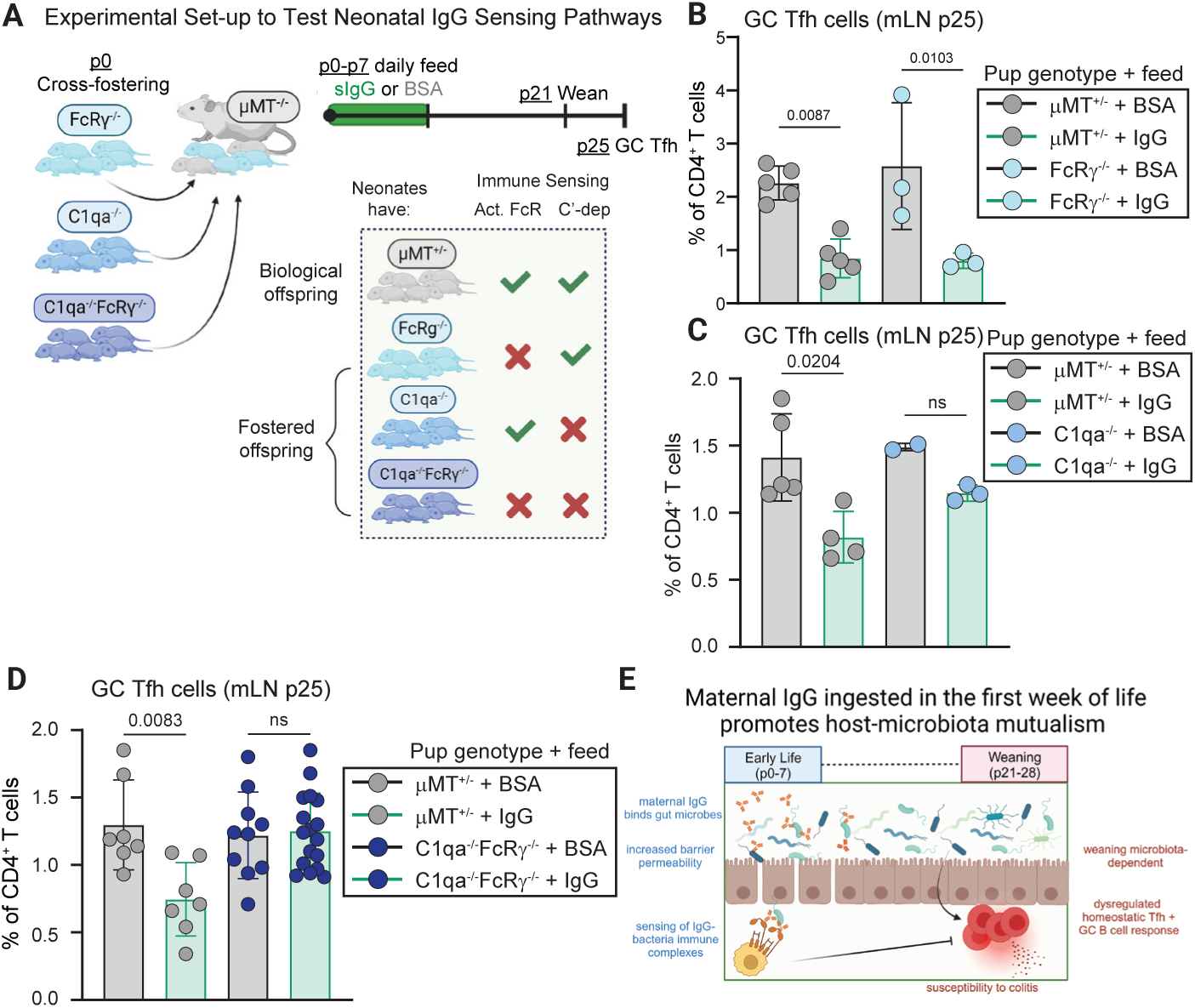
Engagement of Fc-dependent effector functions in the neonate is required for oral IgG to regulate intestinal homeostasis. (**A**) Timed breedings were performed with μMT^-/-^ dams and B6 sires and IgG-sensor deficient dams and sires including FcRγ^-/-^, C1qa^-/-^, and C1qa^-/-^FcRγ^-/-^ pairs. At birth, a portion of offspring of μMT^-/-^ dams and B6 sires were replaced with age-matched IgG-sensor offspring. All offspring were fed 10μg purified sIgG or BSA daily in the first week of life. (**B**) Proportions of mLN GC Tfh cells in p25 FcRγ^-/-^ or μMT^+/-^ offspring fed the indicated proteins in the first week of life. (**C**) Similar to (B) comparing C1qa^-/-^ and μMT^+/-^ offspring. (**D**) Similar to (B) comparing C1qa^-/-^FcRγ^-/-^ and μMT^+/-^ offspring. (**E**) Proposed model: Maternal IgG antibodies, orally acquired via breastfeeding in the first week of life, bind to commensal bacteria in the neonatal gut and form immune complexes, which engage neonatal IgG sensors, thus programming homeostatic immune responses to gut bacteria during the weaning transition. For (**B** to **D**) Error bars indicate the mean ±SD; symbols represent individual mice. Data are representative of two to four independent experiments with ≥3 mice per group. Statistical significance was determined using one-way ANOVA and Tukey post-hoc tests.

Since neither IgG-sensing pathway in isolation was necessary for maternal IgG to restrain T-dependent immune dysregulation, we generated C1qa^-/-^FcRγ^-/-^ mice lacking both IgG-sensing pathways. Strikingly, IgG administration to C1qa^-/-^FcRγ^-/-^ offspring fostered to lactating μMT^-/-^ dams was unable to prevent adaptive immune dysregulation in these animals, which phenocopied control μMT^-/-^-born offspring that did not receive IgG (**Fig 5D**, **Fig S8F-G**). Thus, IgG sensing by the neonatal immune system is required for maternal IgG to calibrate offspring responses to the microbiota.

## Discussion

Early life is a period of profound immune development during which the generation of appropriate responses to newly encountered commensal antigens is essential for health. Here, we show that sensing of maternal IgG-microbiota complexes by the neonatal immune system in early postnatal life instructs subsequent microbiota-triggered immunity during weaning, including preventing dysregulated Tfh and GC B cell responses and alleviating disease severity during experimental colitis. Acquisition of maternal IgG during gestation is well-established to provide neonates passive defense against pathogens(14). Additionally, the microbiota can elicit breastmilk IgG that cross-reacts with enteric pathogens and protects offspring from infection (29, 33). Our present work supports a novel function for breastmilk-acquired IgG in programming the neonatal immune system to limit aberrant responses to commensal organisms.

The engagement of immune signaling pathways in mouse offspring, via complement and activating Fc receptors, distinguishes the mechanism of action of breastmilk IgG from other immunoregulatory pathways that promote intestinal homeostasis in early life. Indeed, breastmilk-derived factors including IgA, growth factors such as EGF, and polysaccharides, largely function by regulating infant immune exposure to gut antigens and/or shaping the assembly or function of the microbiota (5, 34, 35). Differential subclass expression and glycosylation patterns enable IgG to mediate diverse impacts on immune function (30); thus, our findings raise the intriguing possibility that mothers may transmit instructions via breastmilk IgG to tune host-microbiota interactions in offspring. Dissecting both the attributes of maternal IgG and the neonatal cell types and signaling pathways involved in sensing these maternal antibodies remain important areas for future study.

The temporal difference between maternal IgG activity in the first week of life, and subsequent impact on intestinal immunity following weaning provides a regulatory mechanism that is compatible with dynamic and unpredictable changes in microbiota assembly that occur during the weaning transition (36–38). Although our work focuses on maternal IgG-mediated regulation of homeostatic responses to the microbiota, this process may also be involved in regulating immunity to other gut antigens, including pathogens or novel dietary proteins that accompany the introduction of complementary food (39). Human breastmilk also contains IgG (40, 41), and we speculate that the absence of these antibodies in non-breastfed infants impairs microbiota-dependent immune education which may underlie the adverse health outcomes associated with this population (42). Elucidating the maternal-offspring interactions that shape early life immune function will further our ability to promote mutualistic responses to beneficial microbes and prevent pathological responses throughout life.

## Materials and Methods

### Mice

Specific pathogen-free (SPF) C57Bl/6J (Jax000664), μMT^-/-^ (B6.129S2*-Ighm^tm1Cgn^/J,* Jax002288), *C1qa^-/-^* (B6(Cg)-*C1qa^tm1d(EUCOMM)Wtsi^*/TennJ, Jax031675), and TCRβδ^-/-^, (B6.129P2-*Tcrb^tm1Mom^ Tcrd^tm1Mom^*/J, Jax002122) mice were purchased from the Jackson Laboratory. *Fcrg^-/-^*mice generated using C57Bl/6 embryonic stem cells by Dr. Takashi Saito were obtained from a material transfer agreement (RIKEN). Germ-free C57Bl/6 mice were purchased from Taconic (Tac-B6) and germ-free Jh^-/-^ mice were provided by Dr. Andrew Macpherson (Bern, Switzerland). All mice were bred and maintained at an American Association for the Accreditation of Laboratory Animal Care (AALAC)-accredited animal facility at the Fred Hutchinson Cancer Center (SPF animals) or UC Berkeley (germ-free animals) and housed in accordance with the procedures outlined in the *Guide for the Care and Use of Laboratory Animals*. All experiments with mice were performed in accordance with the guidelines of the Animal Care and Use Committee at the Fred Hutchinson Cancer Center or UC Berkeley. All mice were maintained under a 12-h light-dark cycle (7 a.m. to 7 p.m.) and given a standard chow diet (Teklad 2918) and neutral water (pH 6.5-7.5) *ad libitum* unless otherwise indicated.

For cohousing experiments, mice were combined at 3–4 weeks of age until breeding age. For breeding of littermate or cohoused dams, females were separated into individual cages and bred for 2–3 nights, dams were then “re-cohoused” for approximately 18 days. Pregnant females were separated and housed individually until parturition. Age matched offspring of both sexes were used in experiments. In some cases, pups were tattooed at birth via 25G needle (Ketchum).

### Tissue processing

Mice were euthanized with CO_2_, and the mLN and PP were collected and placed into 0.5mL cold complete media (RPMI 1640 supplemented with 2mM L-glutamine, 1mM sodium pyruvate, 100U/ml penicillin, 100mg/mL streptomycin; all from Gibco) supplemented with 3% (v/v) Fetal Calf Serum (FCS; Cytiva)) in a 6 well plate. Tissues were digested by adding 5mL complete media (without FCS) containing 0.5mg/mL DNAseI (Sigma-Aldrich) and 1mg/mL Collagenase III (Stem Cell Technologies) to each well and incubating plates for 20min at 37°C. The enzymatic reaction was stopped with the addition of 10 ml cold complete media containing 3% FCS and tissues were passed through 70 μm (mLN) or 100 μm (PP) cell strainers. The rubber end of 1mL syringes were used to gently mash remaining tissue pieces through the filter, after which the filter was rinsed with complete media to enhance cell recovery. Cells were pelleted at 475x*g*, the supernatant was aspirated, and the pellet was resuspended in 1ml (mLN) or 500μl (PP) complete media containing 3% FCS prior to downstream analysis.

### Flow cytometry

For microbiota flow cytometry, intestinal contents were dissected out, passed over a 40um filter, rinsed with sterile PBS, pelleted at 3210x*g* and resuspended at approximately 5 × 10^7^ bacteria/mL in sterile-filtered Bacterial Staining Buffer (BSB; PBS with 1% bovine serum albumin (BSA; Fisher) and 0.05% Azide (Sigma-Aldrich)). 25 μL of diluted serum or breastmilk antibodies was mixed with 25μL microbial suspension in a v-bottom plate and incubated for 30min at 4°C. 100μl of BSB was added to each well, and cells were pelleted by centrifugation at 3210*g* for 5min. Secondary staining with fluorochrome-conjugated or biotinylated antibodies diluted in BSB followed by streptavidin-PECy7 diluted in BSB was performed (20min each at 4°C). Cells were centrifuged at 3210*g* for 5min and resuspended in sterile-filtered PBS with SYBR Green (Invitrogen) and analyzed by FACS. A similar protocol was used for bacterial flow cytometry, except that an overnight culture of *E. coli* was used as a source of microbes, rather than intestinal contents. See Table S1 for a list of antibody clones/dyes and concentrations used.

For flow cytometry of leukocytes, approximately 2-5×10^6^ cells were incubated in 40 μl of fixable viability dye (Invitrogen) diluted in PBS. This and all subsequent steps were performed in the dark at room temperature unless otherwise stated. After 20 minutes, 10 μl of Fc block diluted in FACS buffer (PBS supplemented with 2% (v/v) FCS and 2mM EDTA; Sigma-Aldrich) was added to cells. After 10 minutes, 50 μl of the extracellular antibody stain diluted in FACS buffer was added. After 30min, 100μl FACS buffer was added to each well, cells were centrifuged at 475*g* for 5min, and fixed and permeabilized using the Foxp3/Transcription Factor Staining Buffer Set (ThermoFisher Scientific) for 20min at 4°C. Cells were centrifuged at 685*g* for 5min, and 100μl of the intracellular antibody stain diluted in Perm Wash was added. After 45min at 4°C, cells were washed with 150μl PermWash, centrifuged at 685*g* for 5min, and resuspended in 200 μl of PermWash. Analysis was performed no later than 18 hours post-staining using a BD FACSymphony A5 High-Parameter Cell Analyzer. Beads (UltraComp eBeads, Invitrogen) stained with individual antibodies were used for compensation, except for the viability dye control, wherein compensation was performed with leftover cells incubated with the viability dye alone. See Table S1 for a list of antibody clones/dyes and concentrations used. To obtain cell counts, a separate aliquot of cells was mixed with beads (Accucheck Counting Beads, Invitrogen) and diluted in 4’,6-Diamidino-2-Phenylindole (DAPI, Sigma Aldrich) and analyzed immediately on a BD FACSymphony A5 High-Parameter Cell Analyzer or BD Canto. All data were analyzed in FlowJo, version 10.

### DSS-induced colitis

Offspring were cohoused at p20 and provided drinking water with 2% DSS (w/v) (molecular weight 36-50 kDa; Thermo-Fisher Scientific) for 7 days. Body weight and physical appearance were monitored approximately every other day until time of sacrifice. Mice that appeared moribund or experienced ≥25% weight loss were euthanized.

### Measurement of Intestinal Permeability

Mice were given 100mg/kg FITC-dex (150kDa, Sigma) or 10mg/kg monoclonal IgG2b (Bioxcel) diluted in PBS via pipet feeding (p0-13) or oral gavage (p14-18). As a positive control, some animals were given substances i.v. Blood was isolated by decapitation (p0-14) or cardiac puncture (p14-28) and allowed to clot for 15min at room temperature. Samples were centrifuged at 13,000*g* for 15min and serum collected. For FITC-dex, serum was diluted 1:5 in PBS and measured with excitation and emission wavelengths of 435 and 538nm, respectively, using an Epoch microplate reader (Biotek). Serum IgG2b titers were assessed via ELISA. Fluorescence or IgG2b levels were interpolated from a standard curve generated via nonlinear 4 parametric logistic regression (Prism, Graphpad).

### Mouse antibody feeding experiments

Pups were fed daily between p0-p7 (8 days) of life with up to 10μg purified antibodies diluted in PBS. Pups were administered up to 2μl of diluted antibodies per day via pipet.

### Enzyme-linked immunosorbent assay (ELISA)

Serum and milk were centrifuged 13,000*g* for 10min and supernatant was used to assay immunoglobulin levels by ELISA. Briefly, Nunc Hi Affinity ELISA plates (Thermo Scientific) were coated with isotype-specific antibodies overnight at 4°C. Plates were washed 5X with Wash Buffer (PBS containing 1%BSA (w/v; Fisher) and 0.05% Tween-20 (v/v; Fisher)) and blocked with PBS with 1% BSA (w/v) and 2% goat serum diluted in PBS (Gibco; v/v) for 1hour at 22-25°C. Plates were washed 5X with Wash Buffer, serial dilutions of samples diluted in PBS with 1% BSA were added, and plates were incubated for up to 2hr at 22-25°C. Plates were washed 5X with Wash Buffer and peroxidase conjugated secondary antibodies specific to mouse isotypes and diluted in PBS containing 1% BSA were added. After up to 2hr at 22-25°C, plates were washed 5X with Wash Buffer, and developed using 1-Step Turbo TMB followed by Stop solution (both Thermo Scientific). Absorbance at 450nm was measured on an Epoch microplate reader (Biotek) within 30min of adding Stop solution. See Table S1 for a list of antibody clones and concentrations used. Antibody titres were interpolated from a standard curve generated via nonlinear 4 parametric logistic regression (Prism, Graphpad).

### Antibody purification from serum and milk

For milk collection, dams were separated from pups for 2 hours at 3-18 days post-parturition and anesthetized via nose-cone with isoflurane. Dams were injected intraperitoneally with 2IU oxytocin diluted in 1mL PBS (Thermo-Fisher Scientific). Milk was collected from mammary glands using microhematocrit capillary tubes, transferred to a microfuge tube and stored at −80°C for subsequent analysis.

For serum collection from adult mice, blood was collected via saphenous vein bleed or cardiac puncture of mice at the time of euthanasia. For neonates, blood was collected after decapitation. To collect serum, blood was allowed to clot at room temperature for up to 60min, centrifuged at 13,000*g* for 15min, and the serum supernatant was transferred to a separate microfuge tube and kept at −80°C for subsequent analysis.

Pooled serum or milk antibodies were purified over a Protein A column according to the manufacturer’s instructions (Pierce; Thermo Scientific). For milk antibodies, milk was thawed on ice, diluted 1:2 in PBS, centrifuged at 21000*g* for 15min, and the top (fat) layer was removed by aspiration. The milk supernatant was transferred to a new tube, diluted a subsequent time 1:2 in PBS and centrifuged again at 21000*g* for 15min. The supernatant (minus fat layer) was used for protein A fractionation. To ensure recovery of all IgG isotypes, including IgG3, IgG was eluted via 2 sequential washes in Elution Buffer pH=2.8, followed by 1 wash with Elution Buffer pH=4.5. IgG was neutralized immediately following each elution and all fractions were combined. IgG enriched and depleted protein fractions were concentrated using a 100,000 MWCO protein concentrator (Pierce; Thermo Scientific). Antibody purity and concentration was assessed by subtype specific ELISA (i.e., IgG, IgA, IgM).

### Milk consumption

Pups were fasted for 4 hours by separating them from lactating dams. Pups were then weighed, returned to dams to nurse for one hour, and weighed a second time. Latching was confirmed by visual confirmation and milk consumption was estimated by the difference in weight gain before and after 1 hour of suckling. To maintain pup health and encourage suckling upon return to dam, pups were kept on a heating pad during the fasting period and maintained in a darkened hood.

### Antibiotic administration

Microbiota depletion was performed as described (25). Briefly, animals were gavaged once per day with a cocktail of antibiotics consisting of ampicillin (20mg/kg; Fisher), vancomycin (10mg/kg; Sigma-Aldrich), neomycin (20mg/kg; ThermoFisher), and metronidazole (20mg/kg; Alfa Aesar) between p17-24. Microbiota depletion was confirmed via stool culture on Luria and blood agar plates, as well as visual confirmation of enlarged cecums.

### Immune complex generation and administration

Serum was collected from TCRβδ^-/-^ mice three weeks after i.v. injection of 1×10^6^ *E.coli* (BL21) or PBS, and incubated with BL21 overnight at 4°C at a ratio of 1ul serum per 1.25×10^6^ bacteria in 50μl PBS. Bacteria were washed in BSB to remove unbound antibodies, resuspended at 5×10^6^ cfu/μl, and kept at 4°C for 8 days. Immune complex formation and stability was confirmed using bacterial flow cytometry at the first and last day of treatment. Pups were fed daily between p0-p7 of life with 5×10^7^ bacteria pre-incubated with sera from immune or naïve animals.

### 16S rRNA Gene Sequencing and Microbial Community Analysis

Amplification and MiSeq Illumina Sequencing of the V4 region of the 16S gene was performed by the Alkek Center for Metagenomics and Microbiome Research (CMMR) at Baylor College of Medicine. Reads were demultiplexed, trimmed and quality checked with bbduk (43). Paired-end reads were then merged using bbmap (43) and run through the MaLiAmPi workflow to generate Amplicon Sequence Variants (ASVs) using DADA2 (44). Pplacer (45) was used to estimate taxonomic classification of ASVs using phylogenetic placement. Alpha diversity estimates were generated by MaLiAmPi and Bray-Curtis dissimilarity was calculated using the R package *vegan* (v2.6-4) to determine beta diversity estimates. A pseudocount was applied prior to calculating differences in the relative abundance of taxonomic groups and ASVs with fewer than 10 reads across all samples were removed from the dataset. Differential abundance was determined using DESeq2 (46), and data was normalized using the “poscounts” size estimator.

### Statistics

Statistical significance was determined as indicated in the figure legends with Prism 10 (GraphPad Software Inc). Significance of beta diversity comparisons was determined using PERMANOVA, which was conducted using the ADONIS function in the R package *vegan*.

## Acknowledgements

We thank the Fred Hutch Comparative Medicine Facility for technical support. We thank G. Barton, G. Reiner, A. Stanbery, and L. Kreuk, for providing germ-free sera and technical assistance regarding germ-free experiments. We thank E. Tait-Wojno, O. Harrison, J. Talbot and P. Mitchell for critical reading of the manuscript. We thank Y. Wu, V. O’Brien, and M. Headley for constructive feedback on this project.

## Author contributions

M.K.S. and M.A.K designed the study experiments and wrote the manuscript. M.K.S performed most of the experiments and data analysis. J.S.S., L.J.M., M.E.C., and D.M.R. assisted with experiments. D.M.R. analyzed 16S rRNA amplicon sequencing data. S.G. assisted with murine milk collection.

## Data and Materials Availability

Sequencing data will be made available on SRA upon publication.

---

**Figure S1.**
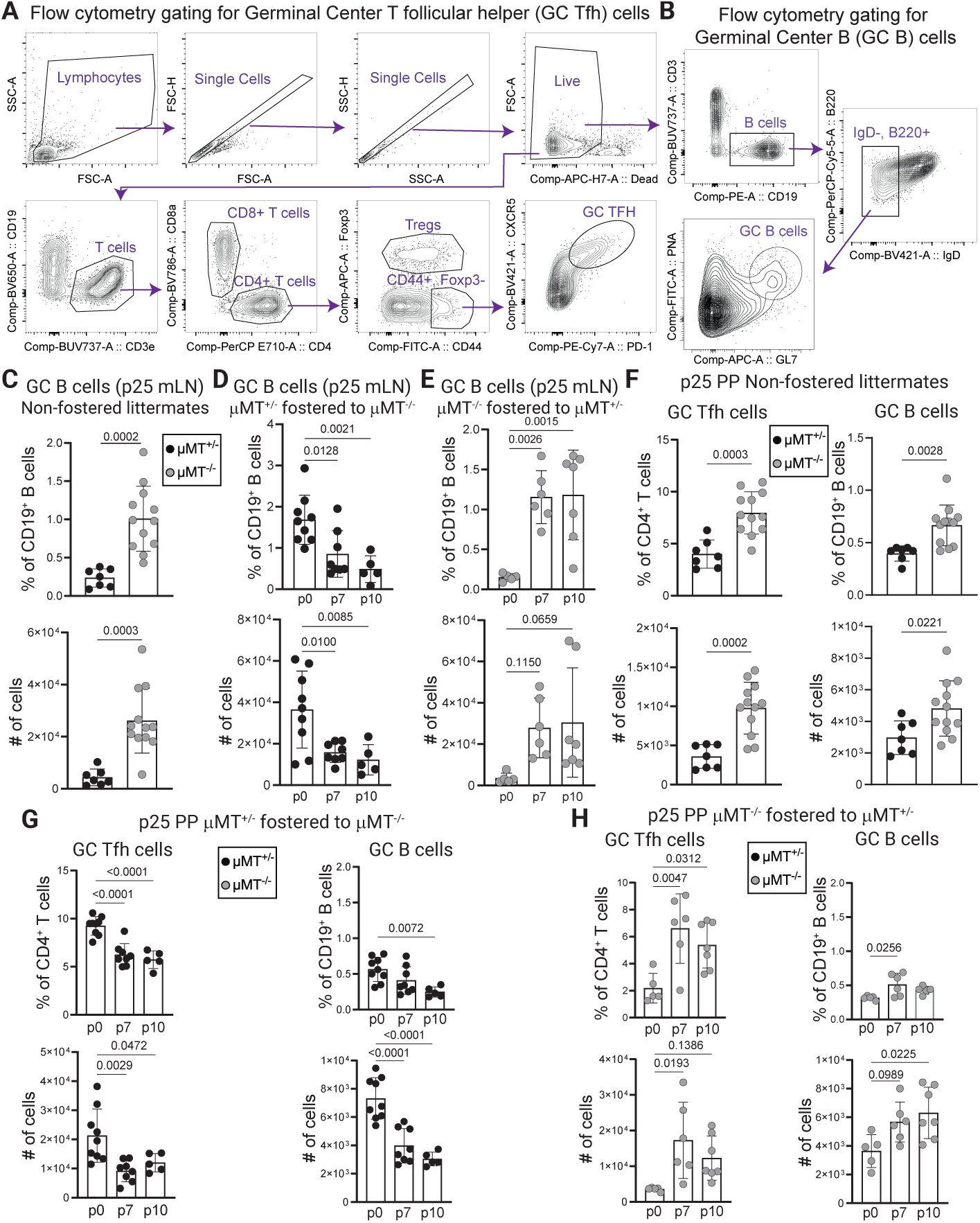
Acquisition of breastmilk antibodies in early life prevents immune dysregulation following weaning. **(A** and **B)** Flow cytometry plots depict the gating scheme for GC Tfh (A) and GC B cells (B) in the mLN and are applicable to other organs including the PP. (**C**) Proportions (top) and numbers (bottom) of mLN GC B cells (CD19^+^B220^+^IgD^-^PNA^+^GL-7^+^) in p25 offspring born to and reared by the indicated dams. (**D**) Proportions (top) and numbers (bottom) of mLN GC B cells in p25 offspring born to μMT^+/-^ dams and fostered to μMT^-/-^ dams at the indicated postnatal timepoints. (**E**) Similar to (D) comparing μMT^-/-^ born offspring fostered to μMT^+/-^ dams. (**F**) Proportions (top) and numbers (bottom) of PP Tfh cells (left) and GC B cells (right) in p25 offspring born to and reared by the indicated dams. (**G**) Proportions (top) and numbers (bottom) of PP Tfh cells (left) and GC B cells (right) in p25 offspring born to μMT^+/-^ dams and fostered to μMT^-/-^ dams at the indicated postnatal timepoints. (**H**) Similar to (G) comparing μMT^-/-^ born offspring fostered to μMT^+/-^ dams. For (**C** to **H**) Error bars indicate the mean ±SD; symbols represent individual mice. Data are representative of four independent experiments with ≥ 5 mice per group. Statistical significance was determined using one-way ANOVA and Tukey post-hoc tests.

**Figure S2.**
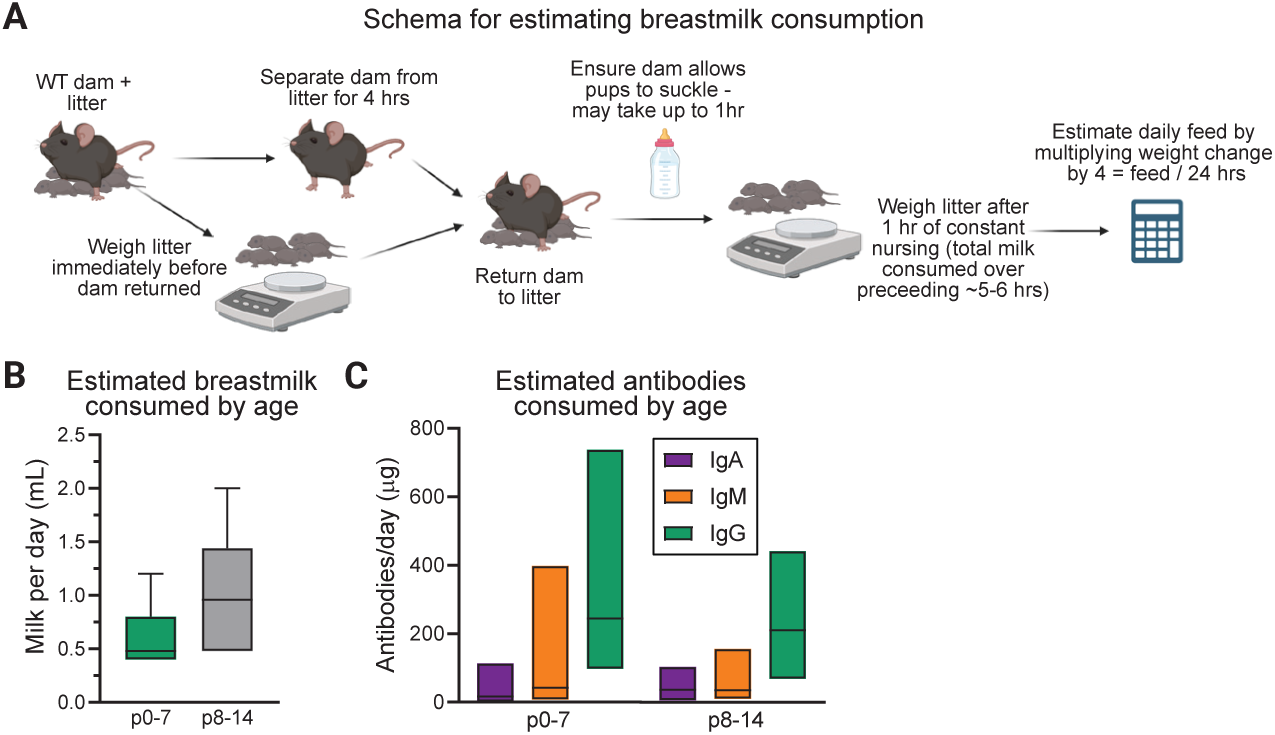
Consumption of breastmilk antibodies restrains microbiota-dependent mucosal immunity in neonates. (**A**) Experimental schema to calculate breastmilk consumption by neonatal mice. (**B**) Estimated daily milk consumption of pups at the indicated postnatal developmental windows. (**C**) Estimated IgG, IgM, and IgA milk antibody consumption of pups as a function of postnatal development. Quantities of consumed antibodies were calculated by multiplying the average volume of milk consumed with the average milk immunoglobulin titer during the same postnatal period. Data are representative of four independent experiments with ≥4 dams per group, ≥4 pups per dam.

**Figure S3.**
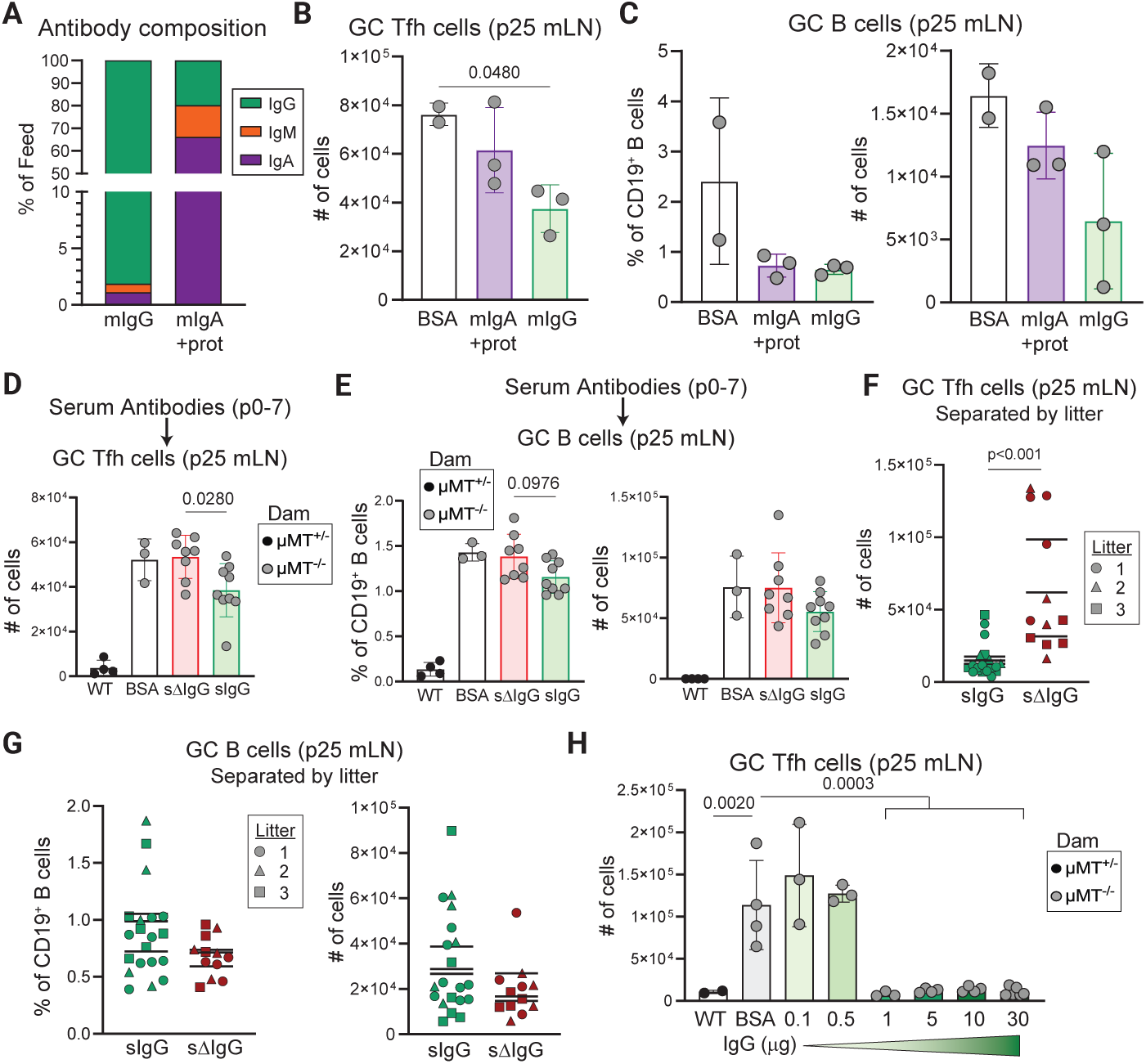
Early life ingestion of IgG suppresses neonatal immune dysregulation. (**A**) Relative amounts of IgA, IgM, and IgG isotypes present in the indicated fractions following Protein A fractionation and concentration of mouse milk. (**B**) Numbers of mLN GC Tfh cells in p25 offspring fed 0.5μg milk (m)IgG, 1μg mIgA (+ other proteins), or 1μg BSA daily from p0-p7, as indicated. (**C**) Similar to (B) comparing proportions (left) and numbers (right) of GC B cells in the mLN of p25 offspring. (**D**) Numbers of mLN GC Tfh cells in p25 offspring fed 10μg serum purified IgG (sIgG), IgG-depleted serum (sdIgG) or BSA daily during p0-p7. (**E**) Similar to (D) comparing proportions (left) and numbers (right) of GC B cells. (**F**) Numbers of mLN GC Tfh cells in littermate, co-housed p25 offspring given 10μg serum IgG or IgG-depleted serum. Littermate animals indicated by symbols. (**G**) Similar to (E) comparing proportions (left) and numbers (right) of GC B cells in the mLN of p25 offspring. (**H**) Numbers of mLN GC Tfh cells in p25 offspring given fed the indicated concentrations of serum IgG or BSA during the first week of life. For (**A**), milk was pooled from approximately two dozen dams, each with ≥4 pups. Pups were between p4-p10. Pooled milk was separated by Protein A chromatography, antibody composition was determined via ELISA, and indicated fractions from pooled milk were used for all independent experiments. For (**B** to **H**) error bars indicate the mean ±SD; symbols represent individual mice. Data are representative of three to four independent experiments with ≥3 mice per group. Statistical significance was determined using one-way ANOVA and Tukey post-hoc tests, with the exception of (**F** and **G**), wherein an unpaired two-tailed Student’s *t* test was used.

**Figure S4.**
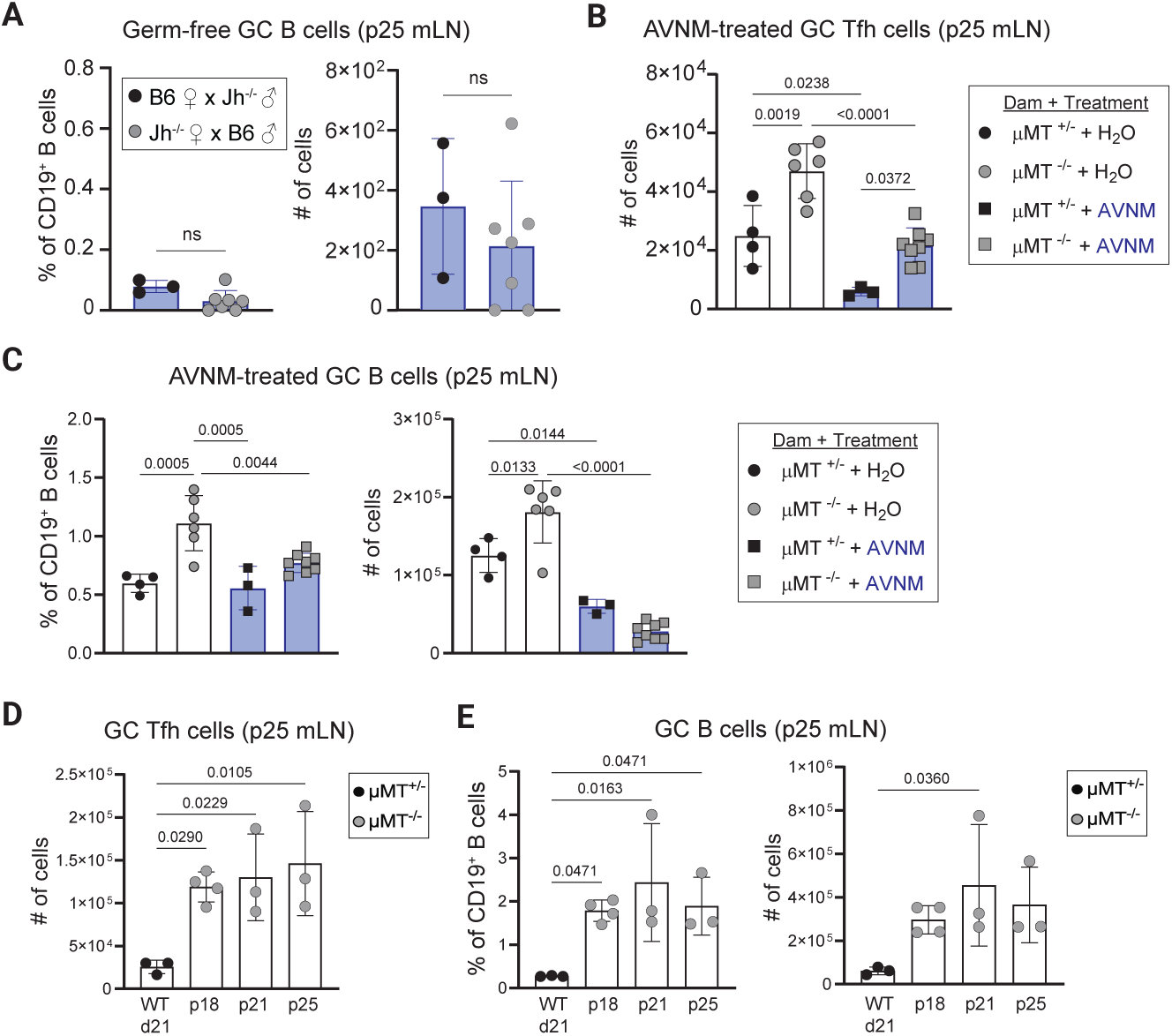
The endogenous microbiota triggers immune dysregulation in the absence of maternal antibodies. (**A**) Proportions (left) and numbers (right) of mLN GC B cells in germ-free p25 offspring born to and reared by the indicated dams bred with the indicated sires. Inset shows cell numbers using a different (lower) scale. (**B**) Numbers of mLN GC Tfh cells in p25 offspring born to the indicated dams and treated with AVNM or water as indicated. (**C**) Similar to (B) comparing proportions (left) and numbers (right) of mLN GC B cells in p25 offspring. (**D**) Numbers of mLN GC Tfh cells in p25 offspring born to the indicated dams and weaned at the indicated postnatal timepoints. (**E**) Similar to (D) comparing proportions (left) and numbers (right) of mLN GC B cells in p25 offspring. Error bars indicate the mean ±SD; symbols represent individual mice. Data are representative of three independent experiments with ≥3 mice per group. Statistical significance was determined using one-way ANOVA and Tukey post-hoc tests, with the exception of (**A**), wherein an unpaired two-tailed Student’s *t* test was used.

**Figure S5.**
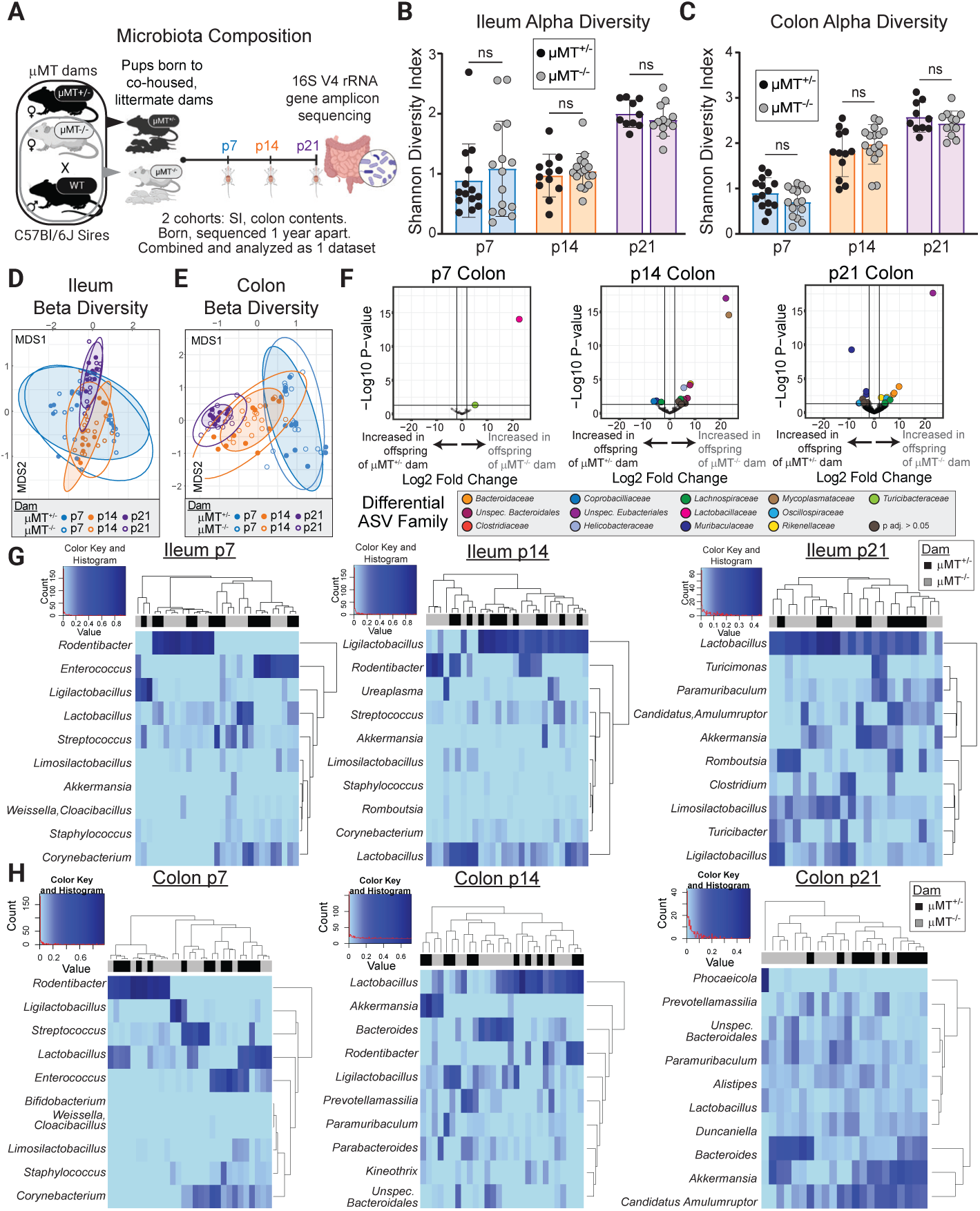
Assembly of the microbiome in the absence of maternal antibodies. (**A**) Timed breedings were performed using co-housed, littermate μMT^+/-^ and μMT^-/-^ dams and B6 sires. Subsets of age-matched litters were euthanized at p7, p14, and p21, and small intestinal and colonic microbiota profiled by 16S rRNA gene sequencing. (**B, C**) Microbial alpha-diversity as calculated by Shannon diversity index of ileal (B), and colonic (C) bacteria isolated from maternal antibody-sufficient and -deficient offspring at the indicated postnatal timepoints. (**D, E**) Non-metric multidimensional scaling plot displaying the Bray-Curtis distances of ileal (D), and colonic (E) bacteria isolated from mice described in (A). (**H**) Number of significant differentially abundant taxa (sequence variants) between maternal antibody-sufficient and -deficient offspring in the indicated intestinal sites at the indicated postnatal timepoints. (**G, H**) Heatmap of genus-level ileal (G) and colonic (H) taxa (rows) isolated from individual offspring (columns) at the indicated postnatal timepoints and ordered by complete linkage clustering. Maternal antibody status of offspring is indicated at the top of each heatmap. PERMANOVA analysis for Maternal antibody status: Ileum p7 p=0.729, F=0.5013; p14 p=0.680, F=0.5131; p21 p=0.416, F=0.9751. Colon p7 p=0.881, F=0.2853; p14 p=0.0.07, F=1.8986; p21 p=0.052, F=1.8883. PERMANOVA analysis for timepoint: Ileum p<0.001, F=20.45. Colon p<0.001, F=14. Data are combined from 2 independent experimental cohorts with samples harvested greater than one year apart totaling n≥10/group. Symbols in (**B** - **E**) and columns in (**G** - **H**) represent individual mice. Statistical significance for (**B** – **C**) was determined unpaired two-tailed Student’s *t* test.

**Figure S6.**
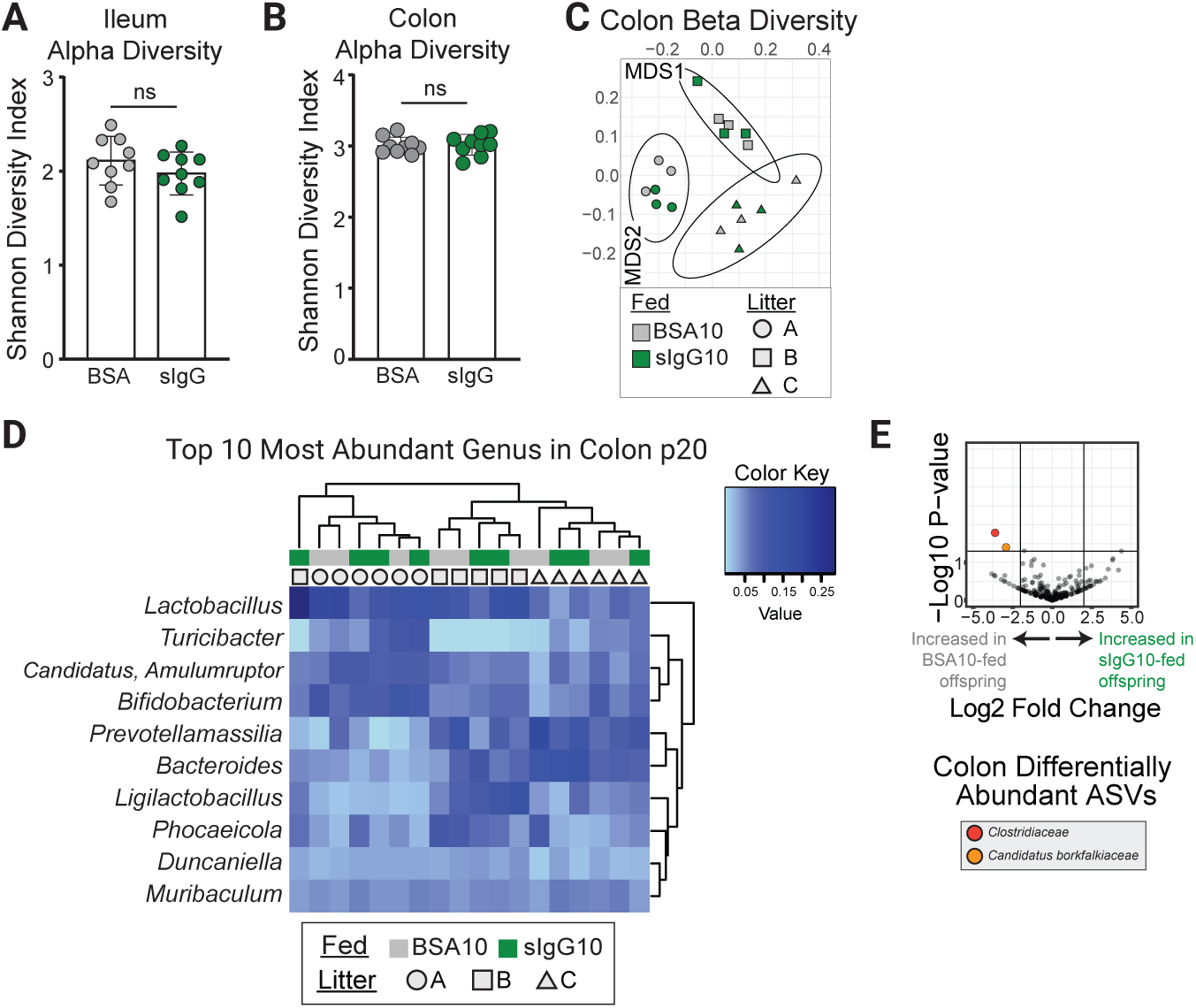
Impact of early life oral IgG in microbiome assembly. (**A, B**) Microbial alpha-diversity as calculated by Shannon diversity index of ileal (A), and colonic (B) bacteria isolated from p20 littermate offspring of μMT^+/-^ dams and B6 sires fed 10 μg sIgG or BSA in the first week of life. (**C**) Non-metric multidimensional scaling plot displaying the Bray-Curtis distances between the 16S profiles of colonic bacteria isolated from mice described in (A-B). PERMANOVA analysis: Litter p=0.001, R^2^= 0.554 and IgG feeding group p=0.917, R^2^=0.02. (**D**) Heatmap of genus-level colonic taxa (rows) isolated from individual offspring (columns) and ordered by complete linkage clustering. Feed and litter status of each mouse is indicated on the top of each heatmap by colors and symbols, respectively. (**E**) Volcano plot of p21 colonic microbiota species (sequence variants). The taxonomic family of significantly altered taxa is indicated. Data are generated from 3 separate litters with n=9 per group. Symbols in (**A** - **C**) and columns in (**D**) represent individual mice. Statistical significance for (**A** – **B**) was determined using an unpaired two-tailed Student’s *t* test.

**Figure S7.**
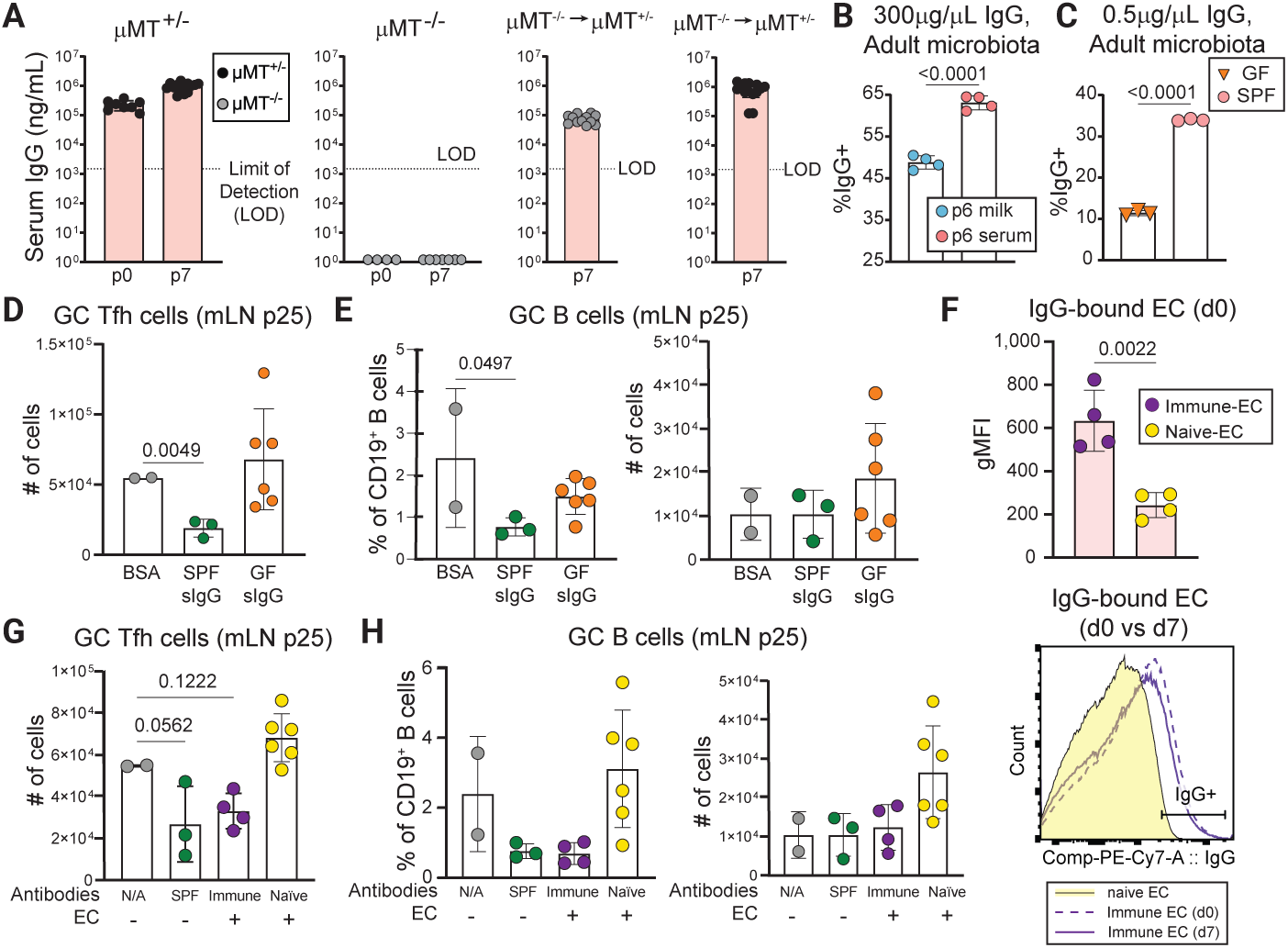
Efficient binding to mucosal antigens is required for early life IgG to restrain mucosal Tfh responses. (**A**) Serum IgG titers in pups born to and/or reared by the indicated dams at p0 and p7. (**B**) mFLOW analysis of the frequency of SYBR+ bacteria bound by IgG purified from the serum and milk of a single donor using intestinal contents isolated from the small intestine and colon of 3-month-old μMT^-/-^ mouse. (**C**) Similar to (B) but comparing IgG-bound bacteria following incubation with sera from SPF or GF animals. (**D**) Numbers of mLN GC Tfh cells in p25 offspring of μMT^-/-^ dams and B6 sires fed 1ug sIgG purified from SPF or GF mice, or BSA, as indicated. (**E**) Similar to (D), comparing proportions (left) and numbers (right) of GC B cells in the mLN of p25 offspring. (**F**) TOP: mFLOW analysis of IgG binding (gMFI) to *E. coli* following incubation with sera from TCRβδo^-/-^ mice challenged i.p. 4 weeks prior with E. coli or PBS, as indicated. BOTTOM: Representative flow plot of data quantified above. (**G**) Numbers of mLN GC Tfh cells in p25 offspring of μMT^-/-^ dams and B6 sires fed 10ug sIgG purified from SPF mice, BSA, or EC immune complexes made using sera from immune or naïve mice, as indicated. (**H**) Similar to (G), comparing proportions (left) and numbers (right) of mLN GC B cells in p25 offspring. Error bars indicate the mean ±SD; symbols represent individual mice. Data are representative of two to four independent experiments with ≥2 mice per group. Statistical significance was determined using one-way ANOVA and Tukey post-hoc tests except for (**B**, **C** and **F**) wherein an unpaired two-tailed Student’s *t* test was used.

**Figure S8.**
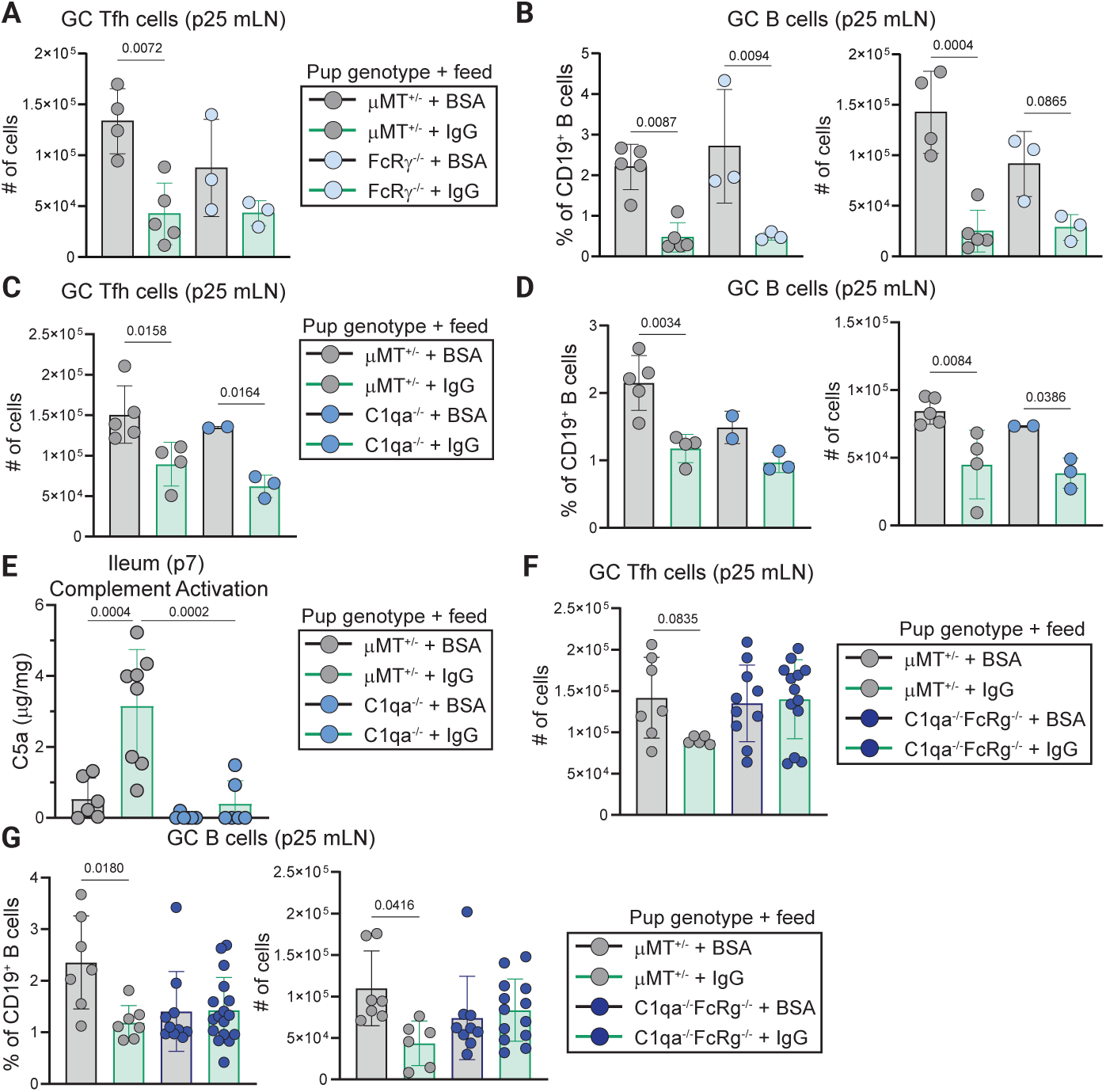
Engagement of Fc-dependent effector functions in the neonate is required for oral IgG to regulate intestinal homeostasis. (**A**) Numbers of mLN GC Tfh cells in p25 FcRγ^-/-^ or μMT^+/-^ offspring fed the indicated proteins in the first week of life. (**B**) Similar to (A) comparing proportions (left) and numbers (right) of mLN GC B cells in p25 offspring. (**C**) Numbers of mLN GC Tfh cells in p25 C1qa^-/-^ or μMT^+/-^ offspring fed the indicated proteins in the first week of life. (**D**) Similar to (C) comparing proportions (left) and numbers (right) of mLN GC B cells in p25 offspring. (**E**) Ileal C5a titers in p7 offspring of μMT^-/-^ dams and B6 sires, or C1qa^-/-^ pairs fed 10μg IgG BSA daily in the first week of life, as indicated. (**F**) Numbers of mLN GC Tfh cells in p25 C1qa^-/-^FcRγ^-/-^ or μMT^+/-^ offspring fed the indicated proteins in the first week of life. (**G**) Similar to (F) comparing proportions (left) and numbers (right) of mLN GC B cells in p25 offspring. Error bars indicate the mean ±SD; symbols represent individual mice. Data are representative of two to four independent experiments with ≥3 mice per group. Statistical significance was determined using one-way ANOVA and Tukey post-hoc tests.

