## Supplemental Figures for "Breastmilk IgG engages the neonatal immune system to instruct host-microbiota mutualism"

Fig.S1

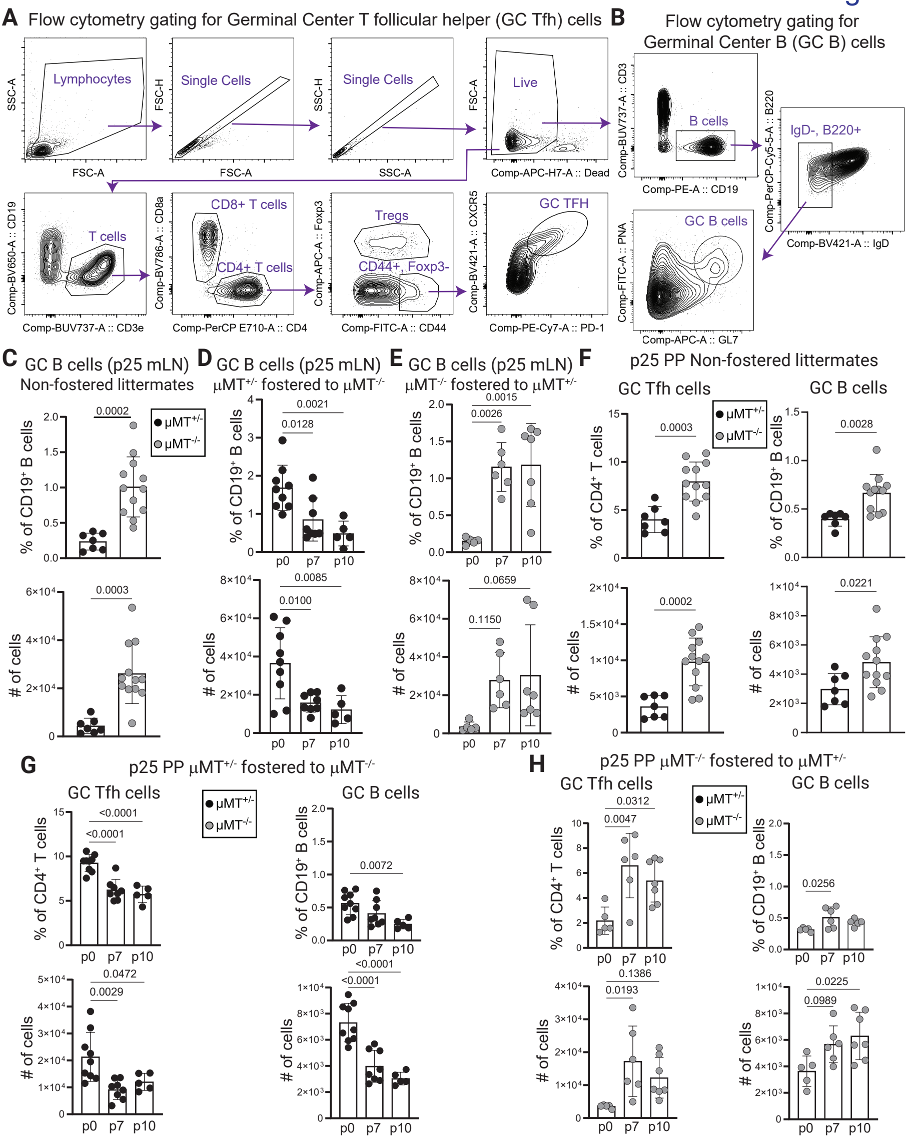

A Flow cytometry gating for Germinal Center T follicular helper (GC Tfh) cells
Lymphocytes
Single Cells
Single Cells
Live
T cells
CD8+ T cells
CD4+ T cells
Tregs
CD44+, Foxp3-
GC TFH
B Flow cytometry gating for Germinal Center B (GC B) cells
B cells
IgD-, B220+
GC B cells
C GC B cells (p25 mLN) Non-fostered littermates
D GC B cells (p25 mLN) μMT+/- fostered to μMT-/-
E GC B cells (p25 mLN) μMT-/- fostered to μMT+/-
F p25 PP Non-fostered littermates
GC Tfh cells
GC B cells
G p25 PP μMT+/- fostered to μMT-/-
H p25 PP μMT-/- fostered to μMT+/-
μMT+/-
μMT-/-
% of CD19+ B cells
% of CD4+ T cells
### of cells
p0
p7
p10

**Figure S1. Acquisition of breastmilk antibodies in early life prevents immune dysregulation following weaning. (A** and **B)** Flow cytometry plots depict the gating scheme for GC Tfh (A) and GC B cells (B) in the mLN and are applicable to other organs including the PP. (**C**) Proportions (top) and numbers (bottom) of mLN GC B cells ($CD19^+B220^+IgD^-PNA^+GL\text{-}7^+$) in p25 offspring born to and reared by the indicated dams. (**D**) Proportions (top) and numbers (bottom) of mLN GC B cells in p25 offspring born to $\mu MT^{+/-}$ dams and fostered to $\mu MT^{-/-}$ dams at the indicated postnatal timepoints. (**E**) Similar to (D) comparing $\mu MT^{-/-}$ born offspring fostered to $\mu MT^{+/-}$ dams. (**F**) Proportions (top) and numbers (bottom) of PP Tfh cells (left) and GC B cells (right) in p25 offspring born to and reared by the indicated dams. (**G**) Proportions (top) and numbers (bottom) of PP Tfh cells (left) and GC B cells (right) in p25 offspring born to $\mu MT^{+/-}$ dams and fostered to $\mu MT^{-/-}$ dams at the indicated postnatal timepoints. (**H**) Similar to (G) comparing $\mu MT^{-/-}$ born offspring fostered to $\mu MT^{+/-}$ dams.

For (**C** to **H**) Error bars indicate the mean ±SD; symbols represent individual mice. Data are representative of four independent experiments with $\geq$ 5 mice per group. Statistical significance was determined using one-way ANOVA and Tukey post-hoc tests.

Fig.S2

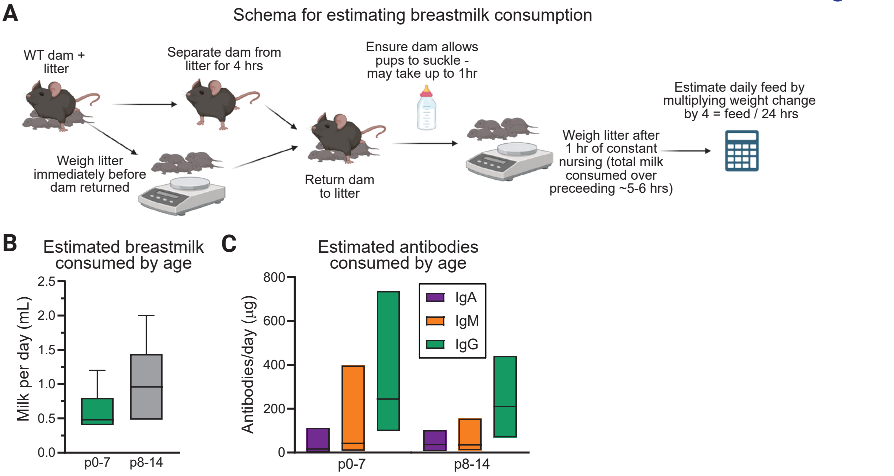
A
Schema for estimating breastmilk consumption
WT dam + litter
Separate dam from litter for 4 hrs
Ensure dam allows pups to suckle - may take up to 1hr
Estimate daily feed by multiplying weight change by 4 = feed / 24 hrs
Weigh litter immediately before dam returned
Return dam to litter
Weigh litter after 1 hr of constant nursing (total milk consumed over preceeding ~5-6 hrs)
B
Estimated breastmilk consumed by age
Milk per day (mL)
2.5
2.0
1.5
1.0
0.5
0.0
p0-7
p8-14
C
Estimated antibodies consumed by age
Antibodies/day (µg)
800
600
400
200
0
IgA
IgM
IgG
p0-7
p8-14

**Figure S2. Consumption of breastmilk antibodies restrains microbiota-dependent mucosal immunity in neonates.** (**A**) Experimental schema to calculate breastmilk consumption by neonatal mice. (**B**) Estimated daily milk consumption of pups at the indicated postnatal developmental windows. (**C**) Estimated IgG, IgM, and IgA milk antibody consumption of pups as a function of postnatal development. Quantities of consumed antibodies were calculated by multiplying the average volume of milk consumed with the average milk immunoglobulin titer during the same postnatal period.

Data are representative of four independent experiments with $\geq$4 dams per group, $\geq$4 pups per dam.

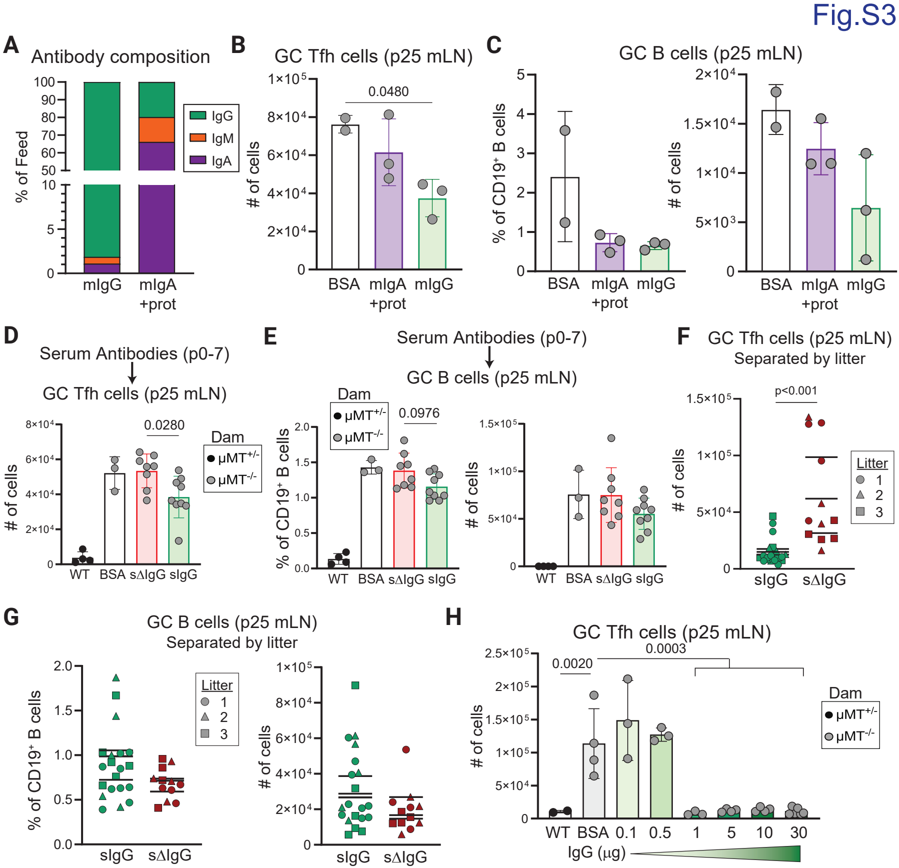

Fig.S3
A
Antibody composition
% of Feed
IgG
IgM
IgA
mIgG
mIgA +prot
B
GC Tfh cells (p25 mLN)
### of cells
0.0480
BSA
mIgA +prot
mIgG
C
GC B cells (p25 mLN)
% of CD19+ B cells
### of cells
D
Serum Antibodies (p0-7)
GC Tfh cells (p25 mLN)
0.0280
Dam
μMT+/-
μMT-/-
WT
BSA
sΔIgG
sIgG
E
Serum Antibodies (p0-7)
GC B cells (p25 mLN)
0.0976
F
GC Tfh cells (p25 mLN)
Separated by litter
p<0.001
Litter
G
GC B cells (p25 mLN)
Separated by litter
H
GC Tfh cells (p25 mLN)
0.0020
0.0003
IgG (μg)

**Figure S3. Early life ingestion of IgG suppresses neonatal immune dysregulation.** (**A**) Relative amounts of IgA, IgM, and IgG isotypes present in the indicated fractions following Protein A fractionation and concentration of mouse milk. (**B**) Numbers of mLN GC Tfh cells in p25 offspring fed 0.5μg milk (m)IgG, 1μg mIgA (+ other proteins), or 1μg BSA daily from p0-p7, as indicated. (**C**) Similar to (B) comparing proportions (left) and numbers (right) of GC B cells in the mLN of p25 offspring. (**D**) Numbers of mLN GC Tfh cells in p25 offspring fed 10μg serum purified IgG (sIgG), IgG-depleted serum (sdIgG) or BSA daily during p0-p7. (**E**) Similar to (D) comparing proportions (left) and numbers (right) of GC B cells. (**F**) Numbers of mLN GC Tfh cells in littermate, co-housed p25 offspring given 10μg serum IgG or IgG-depleted serum. Littermate animals indicated by symbols. (**G**) Similar to (E) comparing proportions (left) and numbers (right) of GC B cells in the mLN of p25 offspring. (**H**) Numbers of mLN GC Tfh cells in p25 offspring given fed the indicated concentrations of serum IgG or BSA during the first week of life.

For (**A**), milk was pooled from approximately two dozen dams, each with ≥4 pups. Pups were between p4-p10. Pooled milk was separated by Protein A chromatography, antibody composition was determined via ELISA, and indicated fractions from pooled milk were used for all independent experiments. For (**B** to **H**) error bars indicate the mean ±SD; symbols represent individual mice. Data are representative of three to four independent experiments with ≥3 mice per group. Statistical significance was determined using one-way ANOVA and Tukey post-hoc tests, with the exception of (**F** and **G**), wherein an unpaired two-tailed Student's *t* test was used.

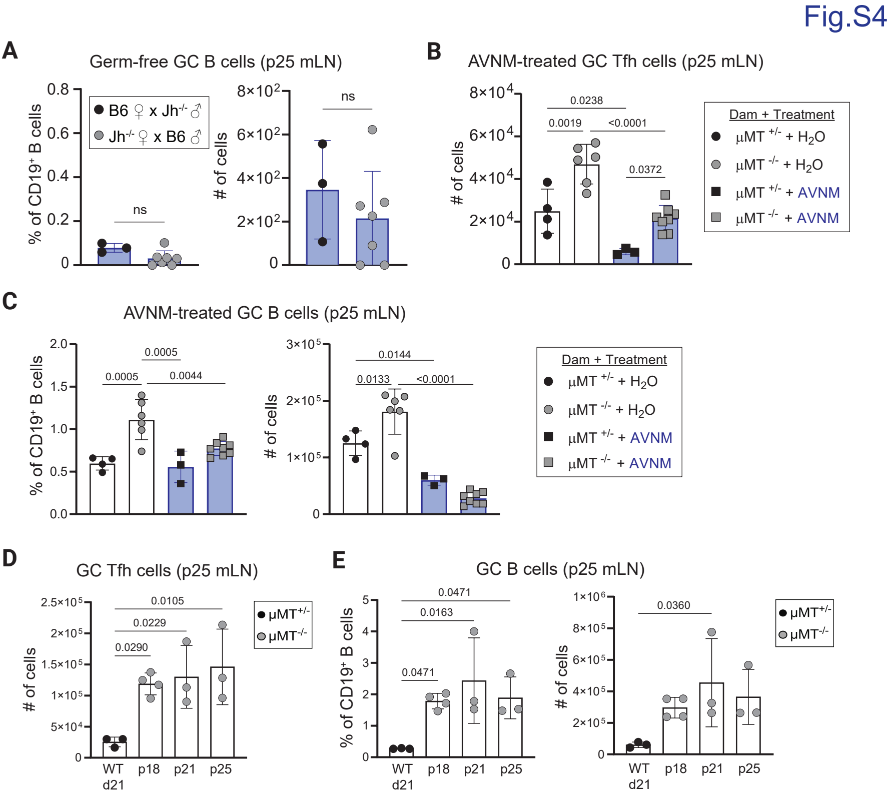

Fig.S4
A
Germ-free GC B cells (p25 mLN)
% of CD19+ B cells
B6 ♀ x Jh-/- ♂
Jh-/- ♀ x B6 ♂
ns
### of cells
ns
B
AVNM-treated GC Tfh cells (p25 mLN)
### of cells
0.0238
0.0019
<0.0001
0.0372
Dam + Treatment
μMT +/- + H2O
μMT -/- + H2O
μMT +/- + AVNM
μMT -/- + AVNM
C
AVNM-treated GC B cells (p25 mLN)
% of CD19+ B cells
0.0005
0.0005
0.0044
### of cells
0.0144
0.0133
<0.0001
Dam + Treatment
μMT +/- + H2O
μMT -/- + H2O
μMT +/- + AVNM
μMT -/- + AVNM
D
GC Tfh cells (p25 mLN)
### of cells
0.0105
0.0229
0.0290
μMT+/-
μMT-/-
WT d21
p18
p21
p25
E
GC B cells (p25 mLN)
% of CD19+ B cells
0.0471
0.0163
0.0471
### of cells
0.0360
μMT+/-
μMT-/-
WT d21
p18
p21
p25

**Figure S4. The endogenous microbiota triggers immune dysregulation in the absence of maternal antibodies** (**A**) Proportions (left) and numbers (right) of mLN GC B cells in germ-free p25 offspring born to and reared by the indicated dams bred with the indicated sires. Inset shows cell numbers using a different (lower) scale. (**B**) Numbers of mLN GC Tfh cells in p25 offspring born to the indicated dams and treated with AVNM or water as indicated. (**C**) Similar to (B) comparing proportions (left) and numbers (right) of mLN GC B cells in p25 offspring. (**D**) Numbers of mLN GC Tfh cells in p25 offspring born to the indicated dams and weaned at the indicated postnatal timepoints. (**E**) Similar to (D) comparing proportions (left) and numbers (right) of mLN GC B cells in p25 offspring.

Error bars indicate the mean ±SD; symbols represent individual mice. Data are representative of three independent experiments with $\geq$3 mice per group. Statistical significance was determined using one-way ANOVA and Tukey post-hoc tests, with the exception of (**A**), wherein an unpaired two-tailed Student's *t* test was used.

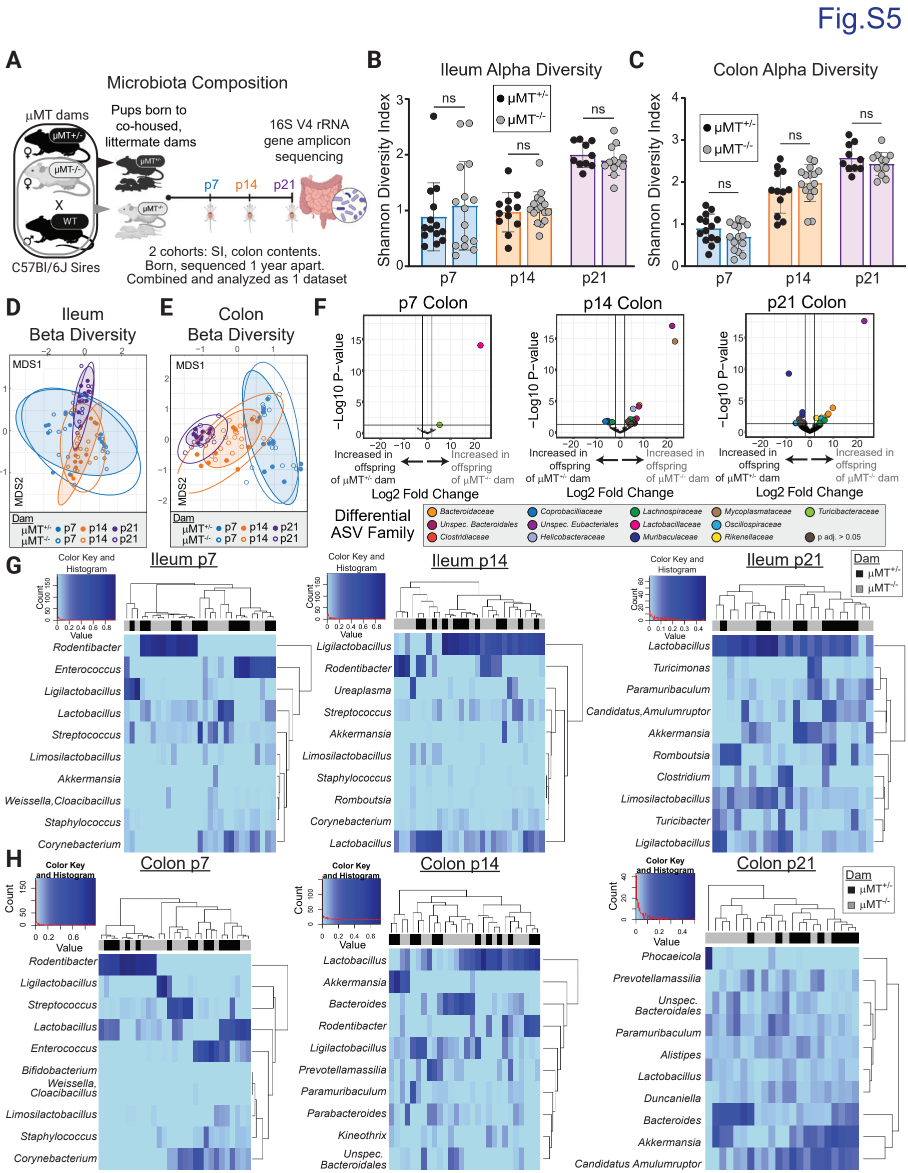
Fig.S5
A
Microbiota Composition
µMT dams
µMT+/-
µMT-/-
X
WT
C57Bl/6J Sires
Pups born to co-housed, littermate dams
16S V4 rRNA gene amplicon sequencing
p7
p14
p21
2 cohorts: SI, colon contents.
Born, sequenced 1 year apart.
Combined and analyzed as 1 dataset
B
Ileum Alpha Diversity
C
Colon Alpha Diversity
Shannon Diversity Index
ns
µMT+/-
µMT-/-
D
Ileum Beta Diversity
E
Colon Beta Diversity
MDS1
MDS2
Dam
F
p7 Colon
p14 Colon
p21 Colon
−Log10 P−value
Increased in offspring of µMT+/- dam
Increased in offspring of µMT-/- dam
Log2 Fold Change
Differential ASV Family
Bacteroidaceae
Coprobacilliaceae
Lachnospiraceae
Mycoplasmataceae
Turicibacteraceae
Unspec. Bacteroidales
Unspec. Eubacteriales
Lactobacillaceae
Oscillospiraceae
Clostridiaceae
Helicobacteraceae
Muribaculaceae
Rikenellaceae
p adj. > 0.05
G
Color Key and Histogram
Count
Value
Ileum p7
Rodentibacter
Enterococcus
Ligilactobacillus
Lactobacillus
Streptococcus
Limosilactobacillus
Akkermansia
Weissella,Cloacibacillus
Staphylococcus
Corynebacterium
Ileum p14
Ligilactobacillus
Rodentibacter
Ureaplasma
Streptococcus
Akkermansia
Limosilactobacillus
Staphylococcus
Romboutsia
Corynebacterium
Lactobacillus
Ileum p21
Lactobacillus
Turicimonas
Paramuribaculum
Candidatus,Amulumruptor
Akkermansia
Romboutsia
Clostridium
Limosilactobacillus
Turicibacter
Ligilactobacillus
H
Colon p7
Rodentibacter
Ligilactobacillus
Streptococcus
Lactobacillus
Enterococcus
Bifidobacterium
Weissella, Cloacibacillus
Limosilactobacillus
Staphylococcus
Corynebacterium
Colon p14
Lactobacillus
Akkermansia
Bacteroides
Rodentibacter
Ligilactobacillus
Prevotellamassilia
Paramuribaculum
Parabacteroides
Kineothrix
Unspec. Bacteroidales
Colon p21
Phocaeicola
Prevotellamassilia
Unspec. Bacteroidales
Paramuribaculum
Alistipes
Lactobacillus
Duncaniella
Bacteroides
Akkermansia
Candidatus Amulumruptor

**Figure S5. Assembly of the microbiome in the absence of maternal antibodies.** (**A**) Timed breedings were performed using co-housed, littermate $\mu MT^{+/-}$ and $\mu MT^{-/-}$ dams and B6 sires. Subsets of age-matched litters were euthanized at p7, p14, and p21, and small intestinal and colonic microbiota profiled by 16S rRNA gene sequencing. (**B, C**) Microbial alpha-diversity as calculated by Shannon diversity index of ileal (B), and colonic (C) bacteria isolated from maternal antibody-sufficient and -deficient offspring at the indicated postnatal timepoints. (**D, E**) Non-metric multidimensional scaling plot displaying the Bray-Curtis distances of ileal (D), and colonic (E) bacteria isolated from mice described in (A). (**H**) Number of significant differentially abundant taxa (sequence variants) between maternal antibody-sufficient and -deficient offspring in the indicated intestinal sites at the indicated postnatal timepoints. (**G, H**) Heatmap of genus-level ileal (G) and colonic (H) taxa (rows) isolated from individual offspring (columns) at the indicated postnatal timepoints and ordered by complete linkage clustering. Maternal antibody status of offspring is indicated at the top of each heatmap. PERMANOVA analysis for Maternal antibody status: Ileum p7 p=0.729, F=0.5013; p14 p=0.680, F=0.5131; p21 p=0.416, F=0.9751. Colon p7 p=0.881, F=0.2853; p14 p=0.0.07, F=1.8986; p21 p=0.052, F=1.8883. PERMANOVA analysis for timepoint: Ileum p<0.001, F=20.45. Colon p<0.001, F=14.
Data are combined from 2 independent experimental cohorts with samples harvested greater than one year apart totaling n$\geq$10/group. Symbols in (**B - E**) and columns in (**G - H**) represent individual mice. Statistical significance for (**B – C**) was determined unpaired two-tailed Student's *t* test.

Fig.S6

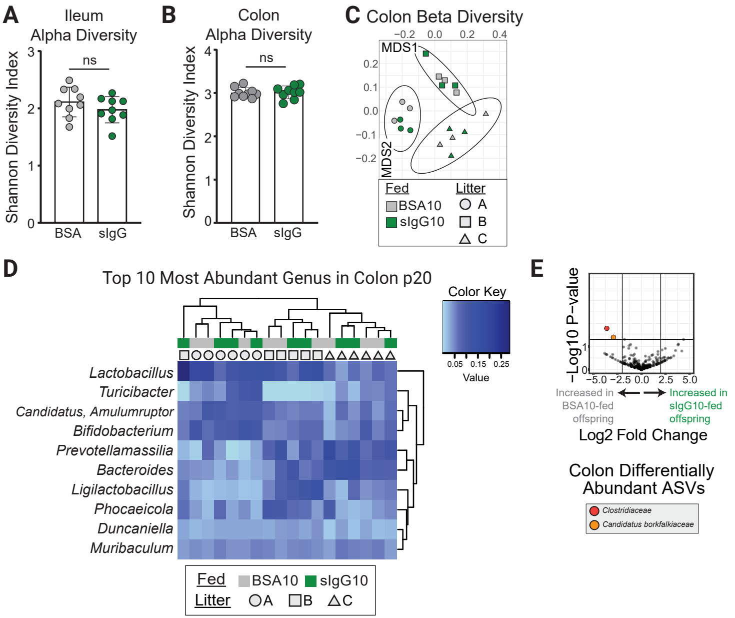

A
Ileum Alpha Diversity
Shannon Diversity Index
ns
BSA
sIgG
B
Colon Alpha Diversity
Shannon Diversity Index
ns
BSA
sIgG
C
Colon Beta Diversity
MDS1
MDS2
Fed
BSA10
sIgG10
Litter
A
B
C
D
Top 10 Most Abundant Genus in Colon p20
Color Key
Value
Lactobacillus
Turicibacter
Candidatus, Amulumruptor
Bifidobacterium
Prevotellamassilia
Bacteroides
Ligilactobacillus
Phocaeicola
Duncaniella
Muribaculum
Fed
BSA10
sIgG10
Litter
A
B
C
E
−Log10 P−value
Increased in BSA10-fed offspring
Increased in sIgG10-fed offspring
Log2 Fold Change
Colon Differentially Abundant ASVs
Clostridiaceae
Candidatus borkfalkiaceae

**Figure S6. Impact of early life oral IgG in microbiome assembly.** (**A, B**) Microbial alpha-diversity as calculated by Shannon diversity index of ileal (A), and colonic (B) bacteria isolated from p20 littermate offspring of $\mu MT^{+/-}$ dams and B6 sires fed 10 $\mu$g sIgG or BSA in the first week of life. (**C**) Non-metric multidimensional scaling plot displaying the Bray-Curtis distances between the 16S profiles of colonic bacteria isolated from mice described in (A-B). PERMANOVA analysis: Litter p=0.001, $R^2$= 0.554 and IgG feeding group p=0.917, $R^2$=0.02. (**D**) Heatmap of genus-level colonic taxa (rows) isolated from individual offspring (columns) and ordered by complete linkage clustering. Feed and litter status of each mouse is indicated on the top of each heatmap by colors and symbols, respectively. (**E**) Volcano plot of p21 colonic microbiota species (sequence variants). The taxonomic family of significantly altered taxa is indicated.

Data are generated from 3 separate litters with n=9 per group. Symbols in (**A** - **C**) and columns in (**D**) represent individual mice. Statistical significance for (**A** – **B**) was determined using an unpaired two-tailed Student's *t* test.

Fig.S7

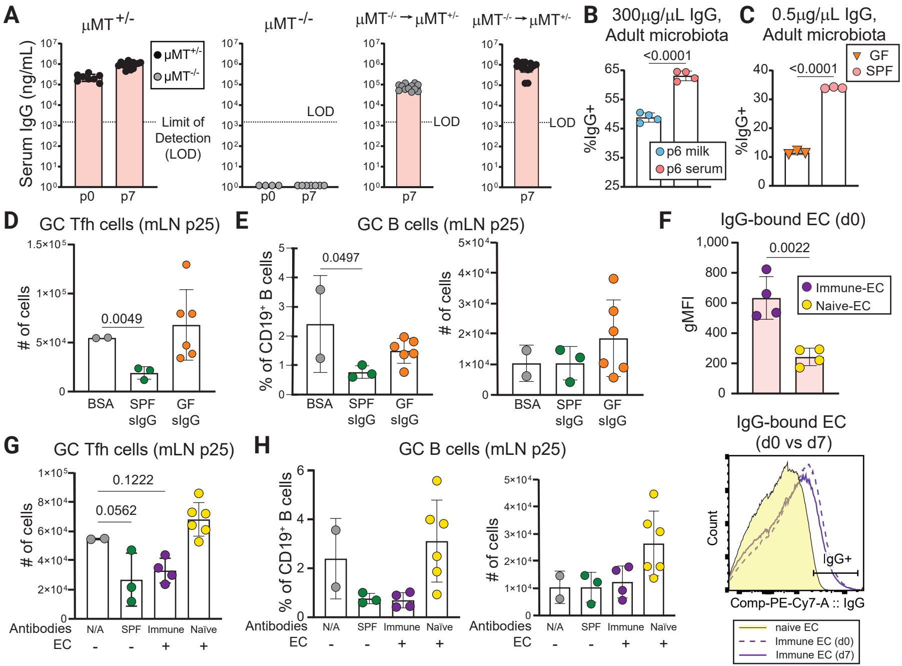
A
μMT+/-
μMT-/-
μMT-/- → μMT+/-
μMT-/- → μMT+/-
Serum IgG (ng/mL)
μMT+/-
μMT-/-
Limit of Detection (LOD)
LOD
p0
p7
B
300μg/μL IgG, Adult microbiota
<0.0001
%IgG+
p6 milk
p6 serum
C
0.5μg/μL IgG, Adult microbiota
<0.0001
GF
SPF
D
GC Tfh cells (mLN p25)
### of cells
0.0049
BSA
SPF sIgG
GF sIgG
E
GC B cells (mLN p25)
% of CD19+ B cells
0.0497
### of cells
F
IgG-bound EC (d0)
0.0022
gMFI
Immune-EC
Naive-EC
G
GC Tfh cells (mLN p25)
0.1222
0.0562
Antibodies
EC
N/A
SPF
Immune
Naïve
H
GC B cells (mLN p25)
IgG-bound EC (d0 vs d7)
Count
IgG+
Comp-PE-Cy7-A :: IgG
naive EC
Immune EC (d0)
Immune EC (d7)

**Figure S7. Efficient binding to mucosal antigens is required for early life IgG to restrain mucosal Tfh responses.** (**A**) Serum IgG titers in pups born to and/or reared by the indicated dams at p0 and p7. (**B**) mFLOW analysis of the frequency of SYBR+ bacteria bound by IgG purified from the serum and milk of a single donor using intestinal contents isolated from the small intestine and colon of 3-month-old $\mu MT^{-/-}$ mouse. (**C**) Similar to (B) but comparing IgG-bound bacteria following incubation with sera from SPF or GF animals. (**D**) Numbers of mLN GC Tfh cells in p25 offspring of $\mu MT^{-/-}$ dams and B6 sires fed 1ug sIgG purified from SPF or GF mice, or BSA, as indicated. (**E**) Similar to (D), comparing proportions (left) and numbers (right) of GC B cells in the mLN of p25 offspring. (**F**) TOP: mFLOW analysis of IgG binding (gMFI) to *E. coli* following incubation with sera from $TCR\beta\delta^{-/-}$ mice challenged i.p. 4 weeks prior with E. coli or PBS, as indicated. BOTTOM: Representative flow plot of data quantified above. (**G**) Numbers of mLN GC Tfh cells in p25 offspring of $\mu MT^{-/-}$ dams and B6 sires fed 10ug sIgG purified from SPF mice, BSA, or EC immune complexes made using sera from immune or naïve mice, as indicated. (**H**) Similar to (G), comparing proportions (left) and numbers (right) of mLN GC B cells in p25 offspring. Error bars indicate the mean ±SD; symbols represent individual mice. Data are representative of two to four independent experiments with $\geq$2 mice per group. Statistical significance was determined using one-way ANOVA and Tukey post-hoc tests except for (**B**, **C** and **F**) wherein an unpaired two-tailed Student's *t* test was used.

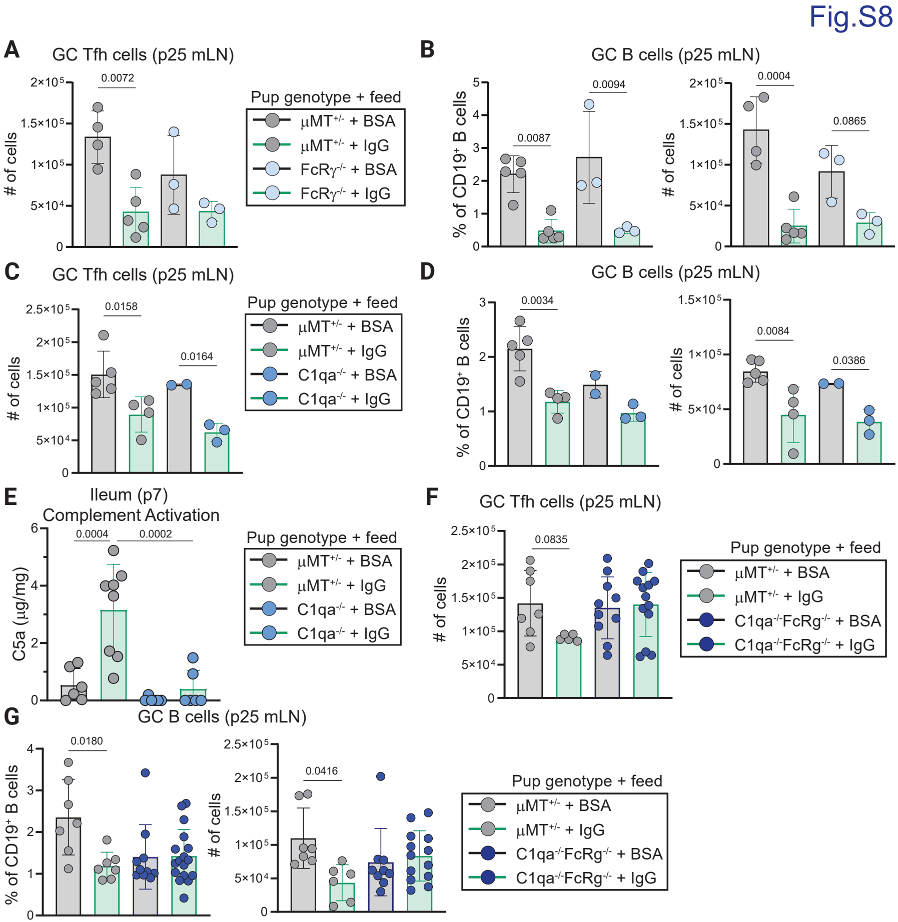
Fig.S8
A
GC Tfh cells (p25 mLN)
### of cells
0.0072
Pup genotype + feed
μMT+/- + BSA
μMT+/- + IgG
FcRγ-/- + BSA
FcRγ-/- + IgG
B
GC B cells (p25 mLN)
% of CD19+ B cells
0.0087
0.0094
### of cells
0.0004
0.0865
C
GC Tfh cells (p25 mLN)
### of cells
0.0158
0.0164
Pup genotype + feed
μMT+/- + BSA
μMT+/- + IgG
C1qa-/- + BSA
C1qa-/- + IgG
D
GC B cells (p25 mLN)
% of CD19+ B cells
0.0034
### of cells
0.0084
0.0386
E
Ileum (p7)
Complement Activation
C5a (μg/mg)
0.0004
0.0002
Pup genotype + feed
μMT+/- + BSA
μMT+/- + IgG
C1qa-/- + BSA
C1qa-/- + IgG
F
GC Tfh cells (p25 mLN)
### of cells
0.0835
Pup genotype + feed
μMT+/- + BSA
μMT+/- + IgG
C1qa-/-FcRg-/- + BSA
C1qa-/-FcRg-/- + IgG
G
GC B cells (p25 mLN)
% of CD19+ B cells
0.0180
### of cells
0.0416
Pup genotype + feed
μMT+/- + BSA
μMT+/- + IgG
C1qa-/-FcRg-/- + BSA
C1qa-/-FcRg-/- + IgG

**Figure S8. Engagement of Fc-dependent effector functions in the neonate is required for oral IgG to regulate intestinal homeostasis.** (**A**) Numbers of mLN GC Tfh cells in p25 FcR$\gamma^{-/-}$ or $\mu$MT$^{+/-}$ offspring fed the indicated proteins in the first week of life. (**B**) Similar to (A) comparing proportions (left) and numbers (right) of mLN GC B cells in p25 offspring. (**C**) Numbers of mLN GC Tfh cells in p25 C1qa$^{-/-}$ or $\mu$MT$^{+/-}$ offspring fed the indicated proteins in the first week of life. (**D**) Similar to (C) comparing proportions (left) and numbers (right) of mLN GC B cells in p25 offspring. (**E**) Ileal C5a titers in p7 offspring of $\mu$MT$^{-/-}$ dams and B6 sires, or C1qa$^{-/-}$ pairs fed 10$\mu$g IgG BSA daily in the first week of life, as indicated. (**F**) Numbers of mLN GC Tfh cells in p25 C1qa$^{-/-}$FcR$\gamma^{-/-}$ or $\mu$MT$^{+/-}$ offspring fed the indicated proteins in the first week of life. (**G**) Similar to (F) comparing proportions (left) and numbers (right) of mLN GC B cells in p25 offspring.

Error bars indicate the mean ±SD; symbols represent individual mice. Data are representative of two to four independent experiments with $\geq$3 mice per group. Statistical significance was determined using one-way ANOVA and Tukey post-hoc tests.
